# Pseudo-repeats in doublecortin make distinct mechanistic contributions to microtubule regulation

**DOI:** 10.1101/808899

**Authors:** Szymon W. Manka, Carolyn A. Moores

**Affiliations:** Institute of Structural and Molecular Biology, Department of Biological Sciences, Birkbeck, University of London, WC1E 7HX, London, UK

**Keywords:** cryo-EM, doublecortin, DCX domain, microtubule-associated protein, pseudo-repeat

## Abstract

Doublecortin (DCX) is a neuronal microtubule-associated protein (MAP) indispensable for brain development. Its flexibly linked DC domains – NDC and CDC – mediate microtubule (MT) nucleation and stabilisation, but it is unclear how. Using high-resolution time-resolved cryo-EM, we mapped NDC and CDC interactions with tubulin at different MT polymerisation stages and studied their functional effects on MT dynamics using TIRF microscopy. Although coupled, each DC repeat appears to have a distinct role in MT nucleation and stabilisation by DCX: CDC is a conformationally plastic tubulin binding module that appears to facilitate MT nucleation by binding tubulin oligomers and stabilising tubulin-tubulin contacts in the nascent MT lattice, while NDC appears to be favoured along the mature lattice, providing enhanced and durable MT stabilisation. Our near-atomic resolution structures of MT-bound DC domains also explain in unprecedented detail the DCX mutation-related brain defects observed in the clinic. This modular composition of DCX reflects a common design principle among MAPs where pseudo-repeats of tubulin/MT binding elements chaperone or stabilise distinct conformational transitions to regulate distinct stages of MT dynamic instability.

## Introduction

The highly dynamic and regulated network of microtubules (MTs) in all eukaryotic cells plays numerous critical roles throughout development. MT-associated proteins (MAPs) spatially and temporally control the MT cytoskeleton (Atherton et al., 2013), thereby governing cell morphology, polarity, intracellular organisation and cell division. MAPs are thus central to organismal morphogenesis and maturation.

Doublecortin (DCX) is the founding member of the family of MAPs that makes significant contributions to MT regulation in metazoan development (Bechstedt et al., 2010; Fourniol et al., 2013; Gönczy et al., 2001). In humans, DCX is indispensable for brain development and specifically for migration of immature neurons, such that lack of functional DCX manifests in grey matter heterotopia or lissencephaly (smooth brain), causing severe intellectual disability and epilepsy (Gleeson et al., 1998; des Portes et al., 1998). Although DCX is predominantly expressed in developing brain, it is also a marker of adult neurogenesis (Moreno-Jiménez et al., 2019), and in several cancers, where its abnormal expression may promote metastasis by facilitating cell migration (Ayanlaja et al., 2017).

MTs are built of α/β-tubulin heterodimers that assemble longitudinally into protofilaments (PFs) and laterally into hollow cylinders. *In vitro*, MTs can form with various PF numbers, but in mammalian cells they mainly contain 13 PFs (Chaaban and Brouhard, 2017; Tilney et al., 1973). DCX strongly promotes nucleation and stabilisation of this physiological 13-PF architecture (Moores et al., 2004). The 13-PF lattice is pseudo-helical, with 12 PFs connected by homotypic (α-α and β-β) lateral contacts and one site, called the seam, built from heterotypic (α-β and β-α) contacts. The molecular mechanisms by which MTs are nucleated, grow with specific architectures and shrink, have been extensively studied but are still not completely understood.

Like numerous other MAPs, DCX is built from pseudo-repeats of MT/tubulin binding elements. It contains two homologous globular doublecortin (DC) domains - NDC and CDC - connected by a 40-residue unstructured linker (Cierpicki et al., 2006; Kim et al., 2003), followed by a disordered C-terminal serine/proline-rich domain (Fig. 1A-C, Supplementary Figs. 1 and 2). NDC and CDC share a similar ubiquitin-like fold but differ in amino acid sequence (24.1% sequence identity) (Fig. 1B). Together, the DC domains have been implicated in MT binding and many disease-causing mutations cluster within them (Bahi-Buisson et al., 2013; Sapir et al., 2000; Taylor et al., 2000) (Supplementary Fig. 2A), but whether the sequence differences between them reflect distinct contributions to DCX function is less clear. An obvious difference between the two DC domains is the length of loop 3, which is 3 residues shorter in CDC (Fig. 1B), while structural studies of individual domains suggest that the biological properties of NDC and CDC are distinct (Burger et al., 2016; Cierpicki et al., 2006; Kim et al., 2003; Rufer et al., 2018). Isolated DC domains alone do not stimulate MT polymerisation, i.e. a DC domain tandem is required for this activity (Taylor et al., 2000), and a distinct contribution of each DC domain to MT lattice-based function of DCX has been indicated (Bechstedt and Brouhard, 2012; Bechstedt et al., 2014; Burger et al., 2016; Liu et al., 2012), although the mechanism by which this occurs is also not known.

**Figure 1.**
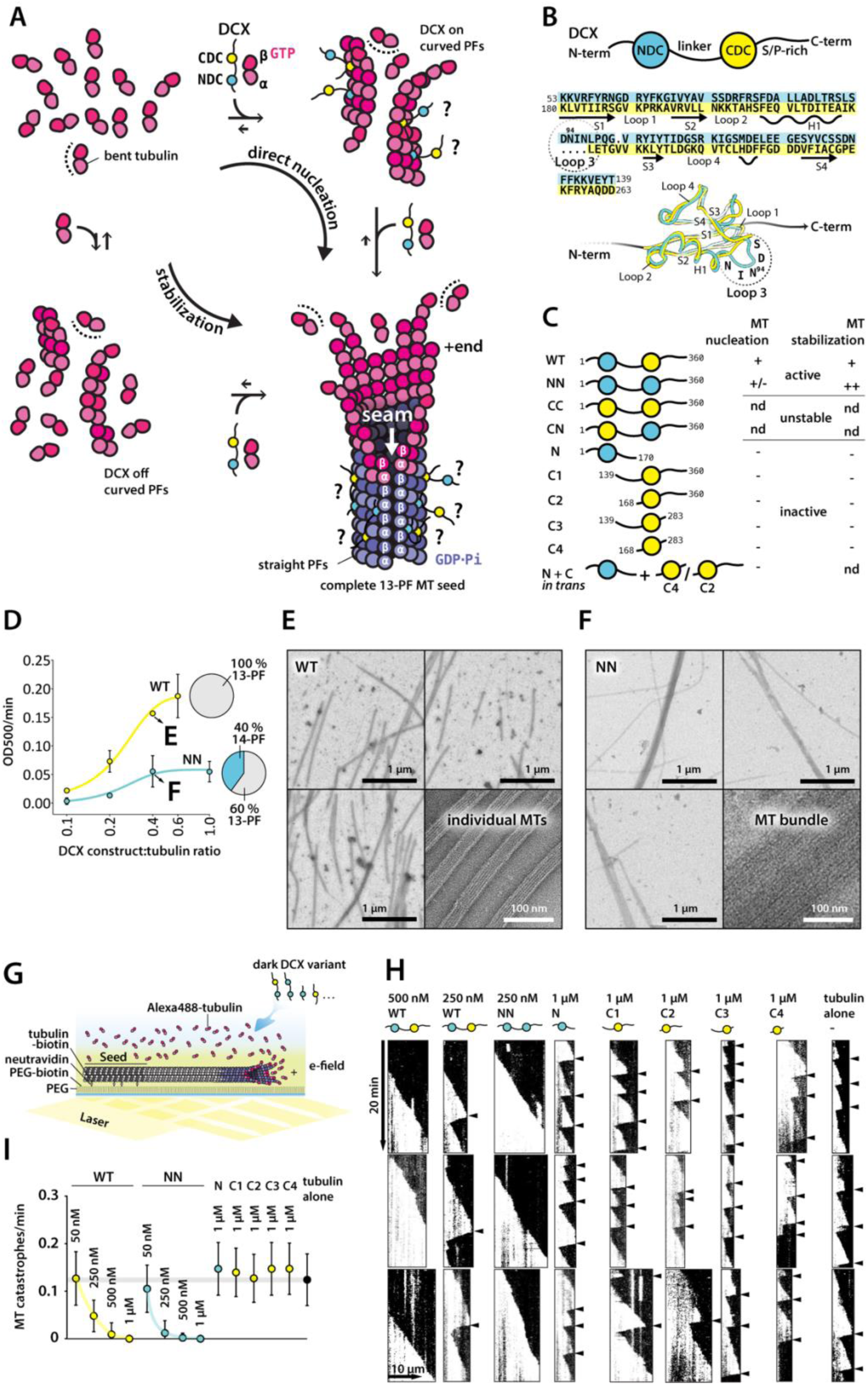
Each DC domain in DCX contributes a distinct function in regulation of MT assembly. **A.** Two routes of MT polymerisation in the presence of DCX are considered: direct nucleation, involving stabilization of early tubulin oligomers by DCX; stabilization, involving DCX binding only between straight PFs of a complete MT lattice. ?, unknown mode of DCX binding to tubulin/MT. **B.** Human DCX schematic, sequence alignment of the DC domains from human isoform 2 (Uniprot ID: O43602-2) with indication of secondary structures and 3D structure alignment, in which selected amino acid positions are indicated using single-letter codes. Sequence identity is 24.1 % by Clustal 2.1 (Larkin, 2007, Bioinformatics, 23, 2947-2948). H, α-helix; S, β-strand. **C.** List of DCX constructs with their abbreviated names and a summary of their activity. Construct boundaries are indicated with residue numbers. +, WT activity; +/-, impaired activity compared to WT; ++, enhanced activity compared to WT; -, no activity; nd, not determined. **D.** Plot of initial MT polymerization velocities at 37 °C in the presence of increasing concentrations of WT or NN. Tubulin concentration = 5 μM. Each point represents an average of 3 independent measurements of the rate of turbidity increase measured at 500 nm wavelength (OD500/min). Fitted are sigmoidal dose-response curves: Y = Y_B_ + ((Y_T_ – Y_B_)/(1 + 10^Xi – X^)), where Y_B_ and Y_T_ are Y values (OD500/min) at the bottom and top plateaus, respectively, and Xi is the X value at the inflection point. Error bar, SD or smaller than a data point. Representative MT samples from 2:5 μM WT:tubulin or NN:tubulin incubations are shown in E and F, respectively. MT architecture distributions for WT and NN are shown with pie graphs. **E-F.** Representative low and high magnification images of MTs nucleated for 10 min at 37 °C from 5 μM tubulin, in the presence of: 2 μM wild-type DCX (WT) (E) or 2 μM NDC-NDC (NN) (F). **G.** Schematic of TIRF microscopy-based MT stabilization assay reported in H and I. Various concentrations of unlabelled DCX variants were added to 9 μM unlabelled tubulin mixed with 1 μM Alexa488-labelled tubulin in the presence of surface-immobilised Alexa488-labelled GMPCPP MT seeds; e-field, evanescent field. **H.** Representative kymographs with arrowheads indicating MT catastrophes. **I.** Quantification of MT catastrophe frequency in the presence of different DCX constructs observed with MT stabilisation TIRF assay. Plotted are mean value with error bar (SD) and smoothed curves connect mean data values. Total free tubulin concentration was 10 μM in all experiments. The basal (tubulin alone) catastrophe frequency level at this tubulin concentration is indicated with a grey bar. Number of MTs analysed across 2-3 movies for each condition: 1 μM WT, n=26; 0.5 μM WT, n=27; 250 nM WT, n=26; 50 nM WT, n=13; 1 μM NN, n=27; 0.5 μM NN, n=26; 250 nM NN, n=25; 50 nM NN, n=19; N, n=18; C1, n=24; C2, n=22; C3, n=18; C4, n=20; tubulin alone, n=29.

DCX does not bind to unpolymerized tubulin, but stabilises existing and/or nascent MT assemblies (Bechstedt et al., 2014; Moores et al., 2006). It recognises a corner between 4 tubulin dimers via a DC domain, except at the seam (Fourniol et al., 2010) (Fig 1A). Cryo-EM studies of DCX-stabilised MTs only reveal density corresponding to a single DC domain – theoretically this could correspond to NDC, CDC or a mixture of both and, so far, available data have not allowed differentiation between these possibilities (Fig. 1A and Supplementary Fig. 2B).

We hypothesised that each DC domain in DCX makes a distinct contribution to MT nucleation and stabilisation, and that the study of DCX could shed light more generally on the mechanisms of these MT processes. We addressed this idea using biochemistry, TIRF-microscopy and high-resolution time-resolved single particle cryo-EM. Near-atomic resolution structural insight of the wild-type DCX (WT) and a chimeric variant, where CDC is replaced with a second NDC (NN), combined with MT dynamics data, uncovered differential roles of the two DC domains. CDC appears to be primarily implicated in MT nucleation, where it also acts to drive specification of 13-PF MTs, whereas NDC appears to collaborate with CDC along the mature MT lattice, providing durable stabilisation of aging MTs.

## Results

### CDC in DCX accelerates MT nucleation and induces 13-protofilament MT architecture

To dissect the roles of the individual DC domains in MT nucleation and stabilisation we began by preparing a series of single and double-DC domain constructs (Fig. 1C, Supplementary Figs. 1C and 2C). The purity of all constructs was verified by SDS-PAGE (Supplementary Fig. 3A) and their intrinsic thermal stability was tested by thermal shift assay (Supplementary Fig. 3B-C). WT, NN and all the single-DC domain constructs had melting temperatures (T_m_) above 60 °C. The construct with swapped DC domains (CN) produced a less sharp thermal shift, but the apparent T_m_ was ∼60 °C (Supplementary Fig. 3C). In contrast, the CDC-CDC (CC) construct had the lowest apparent thermal stability with T_m_ of approximately 40 °C (Supplementary Fig. 3C), suggesting that it is likely partially unfolded at 37 °C, consistent with earlier NMR data (Kim et al., 2003).

**Figure 2.**
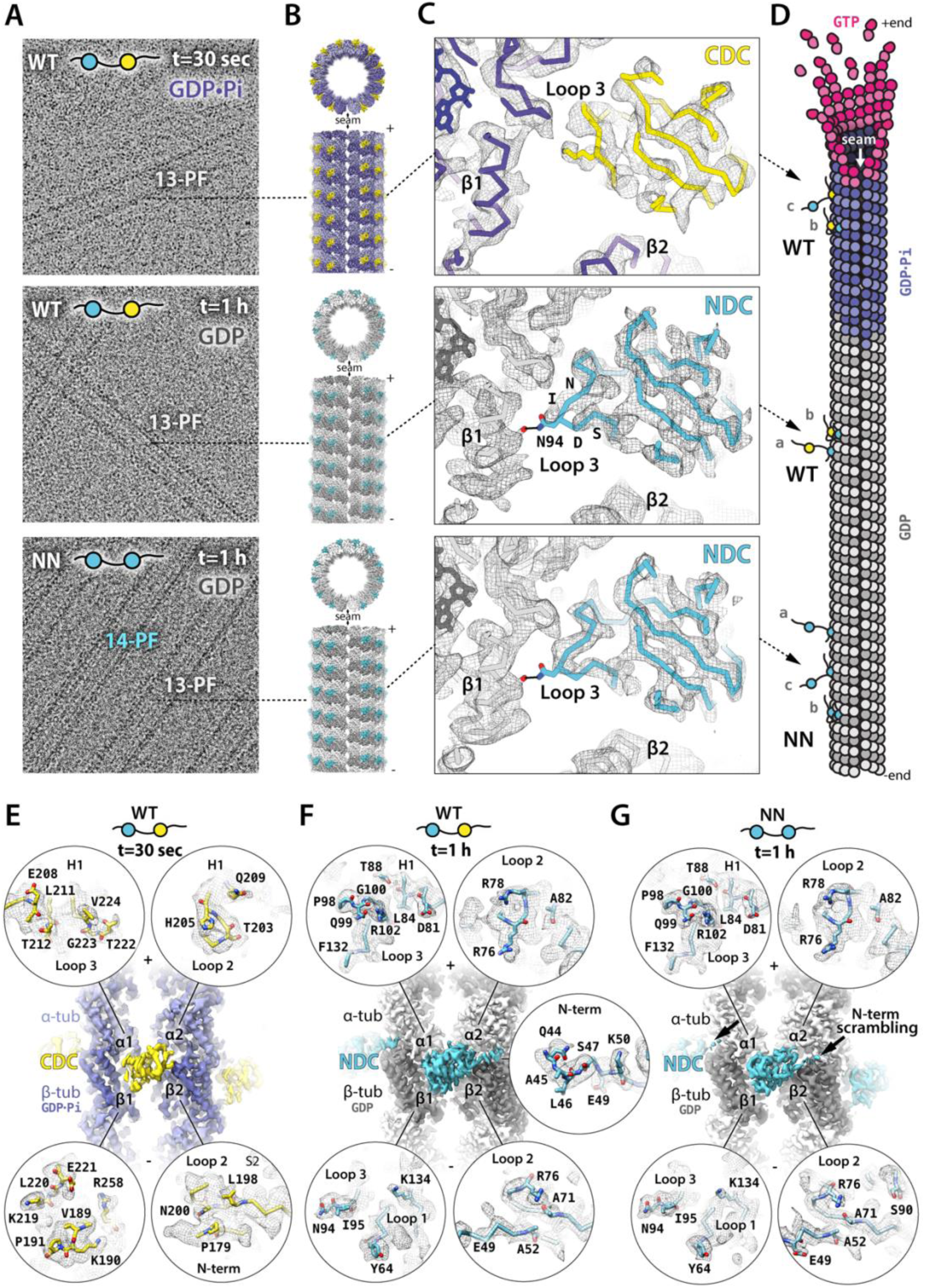
NDC and CDC have distinct roles in stabilising different stages of tubulin assembly. **A**. Representative low-pass filtered cryo-EM micrographs of MTs decorated with wild-type DCX (WT) or NDC-NDC construct (NN) after 30 sec or 1 h incubation at 37 °C. Additional DCX proteins were added in excess after MT deposition on EM grids to maximise MT decoration. The final ratio of WT or NN to tubulin was 50:5 μM, giving dense protein background. WT co-polymerized exclusively 13-PF MTs, while NN both 13- and 14-PF MTs. Rapid (30 sec) MT polymerisation in excess of WT produces MTs in an intermediate GDP.Pi state, whereas long-lived WT-MTs and NN-MTs are all in the GDP state. **B**. Isosurface +end and side views of cryo-EM reconstructions of MTs decorated with different DC domains. α-tubulin, light violet or light grey; β-tubulin, dark violet or dark grey; CDC, yellow; NDC, blue. **C**. Close-up views of the DCX binding site focusing on loop 3 density, which differentiates DC domains. Refined atomic models in predominantly backbone representation are fitted to their respective density maps (wireframe). Selected residues are labelled with single letter codes and selected side chains are shown as sticks. Nucleotide in tubulin subunit designated as β1 is shown as sticks in the top left corner of each panel. Colours as in C, or oxygen atom, red; nitrogen atom, navy blue. **D**. Cartoon illustration of the modes of WT or NN binding to MT lattice, based on cryo-EM results. WT decorates the GDP.Pi lattice predominantly via CDC (c) or via both DC domains (b) and the GDP lattice predominantly via NDC (a) or via both DC domains (b). NN can decorate the GDP lattice via its N-terminal NDC (a), C-terminal NDC (c) or both (b). Modes a and c are compatible with MT cross-linking (bundling) by NN. **E-G**. Isosurface overviews of single DC domains bound to MT lattices together with detailed presentations of all DC domain residues interfacing with MT (4 Å distance), shown in circles representing individual tubulin subunits or binding site of the N-term flanking sequence of WT’s NDC. Only DC residues with at least one atom found within the 4 Å contact zone are fully represented together with their respective densities (wireframe). The density is not masked around the selected residues and occasionally includes adjacent residues that are not displayed in the model. The contact residues are viewed from MT side, i.e. flipped horizontally with respect to the isosurface overviews. Residues upstream of each NDC in NN start to diverge at the point indicated with arrows in G causing scrambling of N-term density. Colouring as in C.

We measured bulk MT nucleation and polymerisation in the presence of our constructs at 37 °C using a turbidity assay, combined with verification of tubulin polymerisation products by negative stain and cryo-electron microscopy (EM) (Fig. 1C-F, Supplementary Fig. 4). We found that only WT and NN efficiently nucleated MTs at 5 µM tubulin concentration - below the ∼10 µM critical concentration (Walker et al., 1991) - and that this activity was dose-dependent (Fig. 1C-F, Supplementary Fig. 4). MTs sporadically occurred at higher concentrations (2 µM) of CN, but products in this reaction were predominantly protein aggregates (Supplementary Fig. 4B). CC induced only protein aggregation (Supplementary Fig. 4B). Thus, both CN and CC were considered unstable and were not analysed further. None of the single-DC domain constructs up to concentrations equimolar to tubulin (5 µM) nucleated MTs, even with both DC domains added *in trans* (Fig. 1C, Supplementary Fig. 4A), suggesting that the physical connection between the DC domains is important. The two longest single-CDC constructs, C1 and C2 (Fig. 1C, Supplementary Fig. 2C) also stimulated tubulin aggregation at high concentrations (3-5 µM) (Supplementary Fig. 4).

Of the tandem DCX constructs that could be analysed further, WT appeared to be a more potent MT nucleator than NN (Fig. 1D). It bundled MTs less than NN (compare Fig. 1E and F), and nucleated solely physiological 13-PF MTs, as expected (Moores et al., 2004). In contrast, ∼40% of MTs nucleated by NN had 14-PF architecture (Fig. 1D and Supplementary Fig. 5A-C). This suggests that CDC in WT not only facilitates MT nucleation, but also participates in defining the resulting MT architecture (Fig. 1A: direct nucleation).

### DC domain tandem is required for MT stabilisation, with CDC not essential for this activity

Bulk MT polymerisation cannot distinguish MT nucleation and stabilisation activities of DCX. To uncouple MT nucleation from stabilisation activity, we used MT seeds stabilised with GMPCPP, a slowly hydrolysable GTP analogue (Hyman et al., 1995). Stabilisation of dynamic MTs grown from these seeds by different DCX constructs was monitored in a total internal reflection fluorescence (TIRF) microscopy assay using fluorescently labelled tubulin (Fig. 1G). Single-DC domain constructs did not reduce MT catastrophe frequency, even at 1 µM concentration (Fig. 1H-I and Supplementary Fig. 5D). Since C1 and C2 showed a tendency to aggregate tubulin (Supplementary Fig. 4), we also tested their MT stabilising capabilities at lower concentrations (250 and 500 nM) and consistently observed no MT stabilisation (Supplementary Fig. 5D). Only WT and NN were capable of preventing MT depolymerisation. NN completely abolished catastrophes at the concentration of 250 nM against 10 µM tubulin, whereas double that concentration (500 nM) of WT was required for the same effect (Fig. 1H-I). At these sub-stoichiometric concentrations both DC domains in principle have unhindered access to their binding sites on the MT lattice. It is thus intriguing that NN has twice as many NDC domains as WT and requires half the concentration of WT to achieve the same MT stabilising effects. This apparent gain of MT stabilisation function upon substitution of NDC for CDC in NN suggests that NDC may have a higher affinity for MT lattice than CDC, and could also explain the potent MT bundling by NN (Fig. 1F).

The ability of the DCX constructs to influence the frequency of MT rescues (resumption of growth after catastrophe) closely followed their ability to influence catastrophe frequency (Supplementary Fig. 5E). MT growth rates were dose-dependently stimulated by NN and C1. They were also significantly increased at higher concentrations (0.5 – 1 µM) of WT, N and C2. This contrasts with what has previously been reported (Bechstedt et al., 2014; Moores et al., 2006) (see Discussion) and suggests that i) at high concentrations, even single-DC constructs are not completely inert regarding MT dynamic instability and ii) the properties of single-DC constructs depend on the presence of other regions from DCX. The MT depolymerisation rates were only significantly slower in the presence of double-DC constructs, WT and NN, as quantified at lower concentrations (50 – 250 nM) where catastrophies still occurred (Supplementary Fig. 5G).

### DCX binds growing MTs initially via CDC, which is subsequently replaced by NDC

Our experiments show that MT nucleation and stabilisation activities of DCX can be differentiated and that each DC domain of DCX is involved in distinct facets of each process. To obtain high-resolution structural evidence of the mode of DCX involvement at different stages of MT assembly we turned to our previous cryo-EM study of MTs polymerised in the presence of WT for 30 seconds or 1 hour. In these experiments, rapid (30 sec) polymerisation captured 13-PF MTs in the transient GDP.Pi state, whereas 1 h polymerisation produced 13-PF MTs in the GDP state (Manka and Moores, 2018). Both samples yielded near-atomic resolution 3D reconstructions of DC domains decorating MT lattice (Fig. 2A-C and E-G, Table 1 and Supplementary Fig. 6), with no density to be assigned to the other regions of the protein, due to their flexibility. Strikingly, the DC domain density in the rapidly polymerised WT-MT reconstruction most resembles CDC, while the corresponding density in the 1 h MT reconstruction resembles NDC (Fig. 2B-C). To verify the conformation of loop 3 in the MT lattice-bound NDC – the main differentiating feature between the DC domains – we carried out an analogous high-resolution 3D cryo-EM study of MTs polymerised for 1 h in the presence of NN. A near-atomic resolution reconstruction of the NN-MT (Fig. 2A-C and G, Table 1 and Supplementary Fig. 6) confirmed the MT lattice-bound structure of NDC observed in the aged WT-MT (Fig. 2C). The NN-MT reconstruction showed somewhat more pronounced density for loop 3 than the WT-MT, suggesting that there could be some mixing of DC domains on the WT lattices (Fig. 2D: a,b). However, overall, these results indicate the involvement of CDC in the early stages of MT assembly, with NDC subsequently joining CDC, and to a large extent replacing it, to facilitate long-term MT stabilisation (Fig. 2D).

**Table 1.**
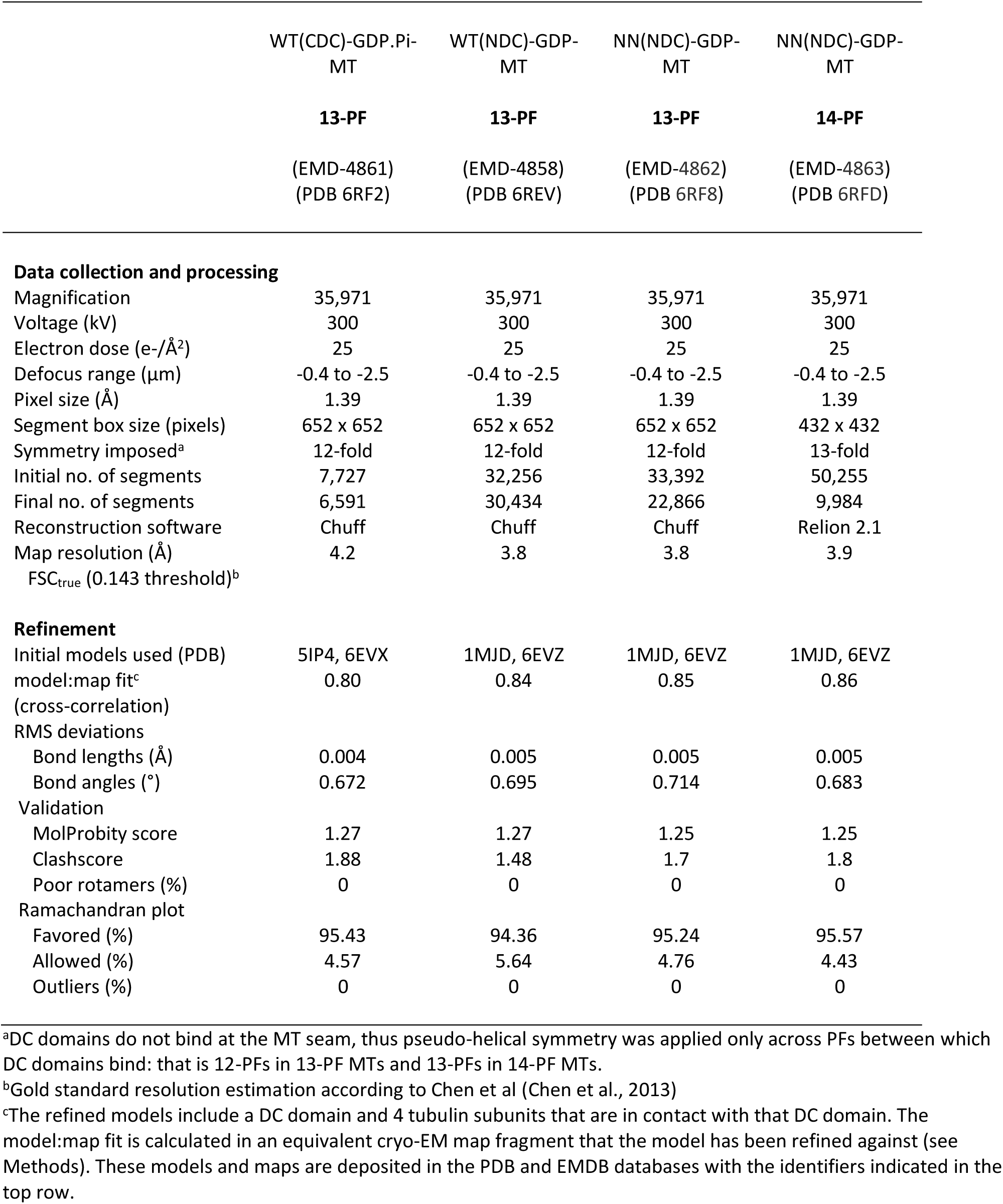
Cryo-EM data collection, atomic model refinement and validation statistics.

### CDC and NDC have distinct binding footprints on the MT lattice

We refined atomic models of CDC and NDC together with the bound tubulin subunits directly in their respective density maps (Fig. 2C and E-G and Table 1) and analysed the CDC-MT and NDC-MT interfaces (defined as residues situated within 4 Å distance from tubulin (Fig. 2E-G). Both CDC and NDC interact with all 4 tubulin subunits at the lattice vertex, but with distinct tubulin residues.Overall, MT interactions by both NDC and CDC are mainly electrostatic, including several hydrogen bonds and/or salt bridges (Supplementary Fig. 7), but also involve hydrophobic contacts. In aged WT-MTs, additional interactions are mediated by the N-terminal region flanking NDC (residues 44-50; Fig. 2F). This likely confers MT lattice affinity enhancement to NDC, supporting the predominance of bound NDC in this reconstruction. No such contact is seen when CDC is bound (Fig. 2E). Furthermore, the equivalent region is poorly represented in NN-MT reconstruction upstream of residue 48, where the domain cloning boundary has been set (Fig. 2G, Supplementary Fig. 2C). The deterioration of density from the point at which sequences flanking each NDC of NN start to diverge implies that it is very likely that either or both of the NDCs bind the MT lattice and are averaged together (Fig. 2D: a,b,c). Otherwise, NDC refined in WT-MT map and that refined in NN-MT map show almost identical 4 Å contacts (Fig. 2F-G), apart from S90 residue of NDC which is detected only in NN-MT reconstruction (Fig. 2G, β2 corner).

Our cryo-EM models suggest how CDC can be replaced by NDC on the MT lattice. Not only does the N-terminal flanking region of NDC make additional contacts with the MT wall compared to CDC (Fig. 2E-F), but also its longer loop 3 is inserted in the inter-PF grove. This loop, by closely following the shape of the MT wall, makes an additional polar contact via N94 with β-tubulin’s D209 and a hydrophobic contact via I95 with β-tubulin’s A302. Altogether, CDC forms 9 and NDC forms 11 ionic interactions with MT lattice (Supplementary Fig. 7), which suggests higher affinity of NDC for MT lattice compared to CDC.

### NDC binding to MT lattice is not selective for a particular MT architecture

Since NN-stimulated MT nucleation produces 14-PF almost as well as 13-PF MTs, we investigated whether there were any architecture-specific differences in NDC observed in these MTs. The manual collection of NN-MT cryo-EM data for high-resolution 3D reconstruction was focused on the physiological 13-PF architecture to enable direct comparisons with WT-MT reconstruction. Nevertheless, supervised 3D classification of all picked NN-MT segments revealed a 13-PF class containing 73 % and a 14-PF class containing 26 % of segments. The remaining 1 % are particles of relatively poor quality. The 14-PF NN-MT reconstruction had a resolution nearly equivalent (3.9 Å) to that of the 13-PF NN-MT reconstruction (3.8 Å) (Fig. 3A-B, Table 1 and Supplementary Fig. 6). Crucially, NDC density is equally pronounced in both the 13-PF and 14-PF NN-MT 3D maps (Fig. 3A-B). Moreover, the level of decoration with NN of the 13/14-PF lattices is comparable to that of 13-PF lattices with WT, as evidenced by analyses of MT Fourier transforms (Fig. 3C, Supplementary Fig. 8). NDC requires only minimal structural adaptation to fit to the slightly narrower inter-PF angle of the 14-PF lattice compared to the 13-PF lattice (Fig. 3D). Overall, NDC appears to be relatively insensitive to the inter-PF angle, with identical (within the resolution) polar NDC-MT interactions formed in each MT architecture (Supplementary Fig. 7).

**Figure 3.**
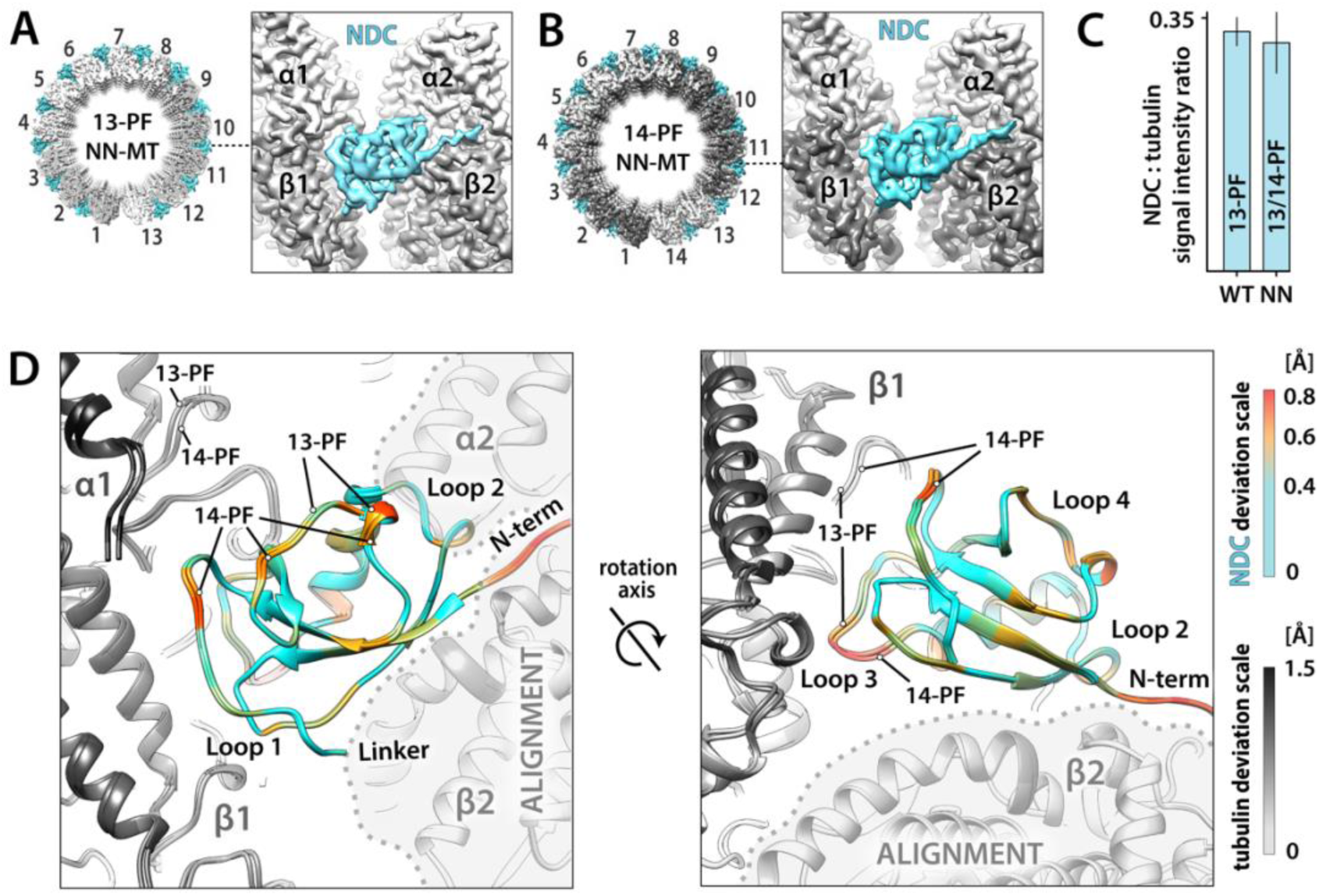
NDC proficiently recognizes and stabilizes both 13-PF and 14-PF MTs. **A**. Isosurface top view of the whole 13-PF MT cylinder and a side view close-up on NN binding site at the vertex of 4 tubulin dimers, where NDC of NN decorates the MT lattice. α-tubulin, white; β-tubulin, grey; NDC, blue. The valley between PFs 1 and 13 represents MT seam, where NN does not bind. **B**. Isosurface views as in A of the 14-PF MT decorated with NN. α-tubulin, light grey; β-tubulin, dark grey; NDC, blue. The valley between PFs 1 and 14 represents MT seam, where NN does not bind. **C**. Relative extent of MT lattice decoration based on average intensities of layer lines representing NDC (8 nm periodicity) and tubulin (4 nm periodicity) in averaged Fourier transforms of WT-MT and NN-MT segments, as described in Supplementary Fig 8 and Methods. Error bar, SD. **D**. Comparison of the DCX binding clefts in the 13-PF and 14-PF MTs and mapping of the associated small (<0.8 Å) conformational adjustments in NDC. NN-MT lattice fragments were aligned on the indicated α2 and β2 tubulin subunits (ALIGNMENT). Deviations in tubulin are depicted with grey scale and in NDC with colour scale. Two perspectives are shown to cover all major MT-binding loops of NDC.

### DC domain plasticity and its potential role in MT assembly

NDC shows structural rigidity regardless of the architecture of the MT to which it is bound, and this MT-bound conformation is highly similar to the published NMR and X-ray structures of NDC (Fig. 4A, cyan, root-mean-square deviation, RMSD = 1.3 Å). In contrast, the conformation of the MT-bound CDC in our reconstruction is substantially different from that stabilized in its crystal structure (Burger et al., 2016) (Fig. 4A, yellow, RMSD = 4.3 Å). We wondered if this apparent structural plasticity of CDC could be related to its preferential association with the transient GDP.Pi-MT state, as revealed in our cryo-EM experiments, as there is no obvious mechanism how the lattice itself could dictate this preference. The GDP.Pi lattice is locally similar to the GDP lattice at the DCX binding site, with the greatest deviation of tubulin backbone between the two states being at the α2 corner of the DC binding site, amounting to ∼1 Å (Manka and Moores, 2018) (Supplementary Fig. 9). It has previously been proposed that DCX can recognise longitudinal curvature of MTs via CDC (Bechstedt et al., 2014), and this could underlie the interaction with early, curved tubulin assemblies (Fig. 1A: direct nucleation), leading to generation of complete MTs for subsequent elongation (MT growth). We therefore wondered if CDC might be specifically involved in stabilising such early intermediates of MT nucleation, due to its conformational flexibility, i.e. binding bent tubulin assemblies via one conformation and the early (GDP.Pi) MT lattice via the other.

**Figure 4.**
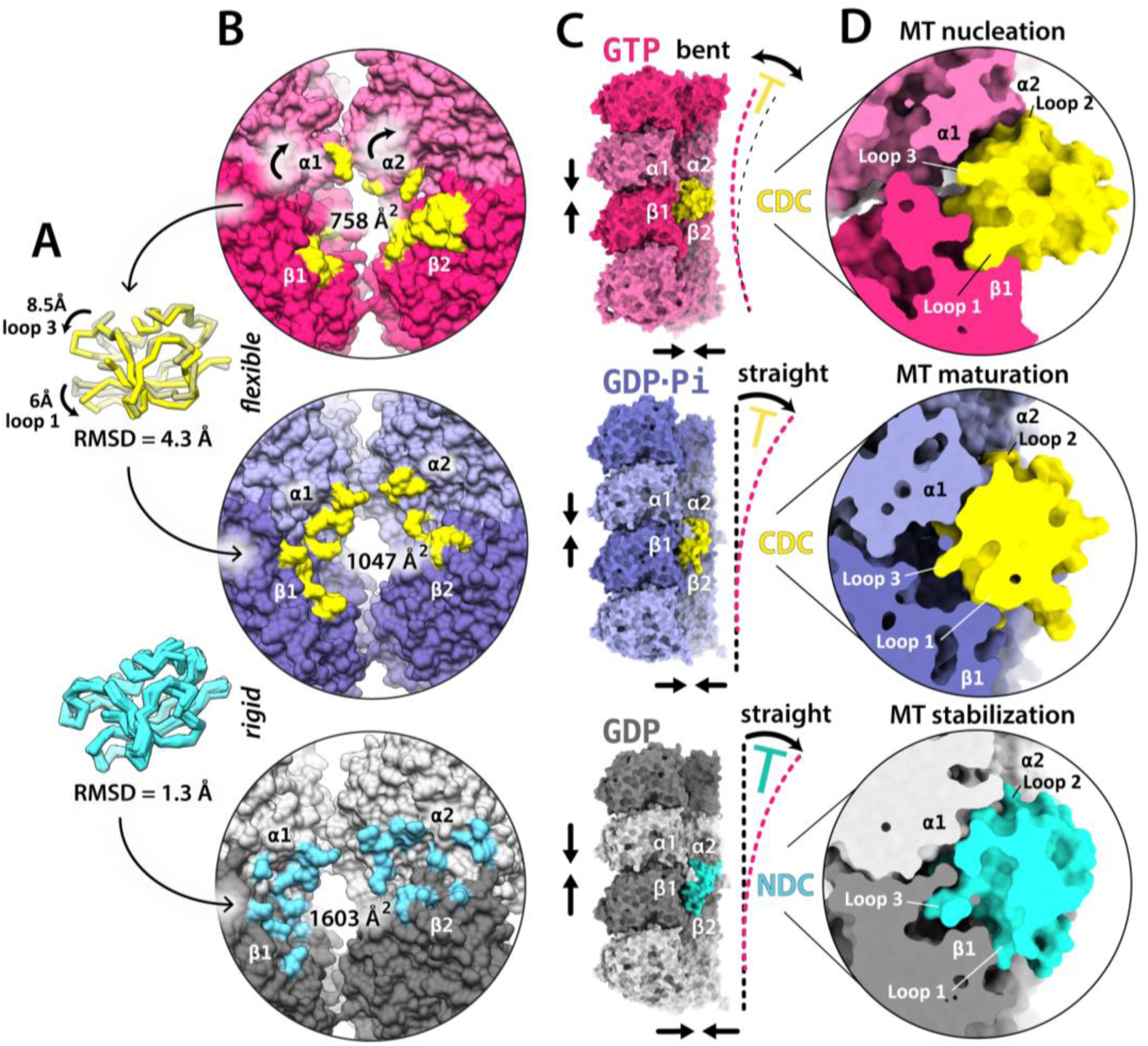
Sequential action of the DC domains according to their tubulin/MT-binding affinity and conformational plasticity. **A**. Superposition of CDC X-ray structure 5IP4 (darkened and transparent) with the GDP.Pi-MT-binding conformation (this study; bright yellow) and superposition of 7 NMR and X-ray NDC structures from Protein Data Bank (PDBs: 5IO9, 1MJD, 1UFO, 2BQQ, 2NDF, 5IKC, 5IN7) with the GDP-MT-binding conformation (this study). Molecular modelling and the intrinsic structural plasticity of CDC predict its ability to interact with bent as well as straight tubulin assemblies by flexing its loops (see also Movie 1), whereas NDC shows conformational stability. **B**. DC domain binding footprints (residues within 4 Å distance) on a bent GTP-tubulin assembly (putative) and an MT lattice at GDP.Pi and GDP states (cryo-EM). Bent arrows, direction of α-tubulin motion during bent-to-straight (GTP to GDP.Pi) tubulin transition (Movie 1). Interface areas were calculated using PDBePISA (http://www.ebi.ac.uk/pdbe/pisa/). **C**. Overview of the putative (modelled) structure of CDC bound at the vertex of 4 bent tubulin dimers (GTP state) and two cryo-EM structures of CDC and NDC bound at the vertex of 4 tubulin dimers in GDP.Pi- and GDP-MT lattice conformation, respectively. Straight arrows indicate stabilization of tubulin-tubulin contacts. The colour-coded T shapes suggest DC domain-dependent inhibition of structural change: CDC binding between bent tubulin dimers may restrict longitudinal curvature of the assembly and the subsequent CDC and NDC binding to MT lattice appears to lock tubulin in the lattice-specific straight conformation. **D**. Close-up cutaway views of the complexes in C.

To test the viability of our hypothesis we turned to molecular modelling using Rosetta (Gray et al., 2003) (Fig. 4B-D and Supplementary Fig. 10), since the non-MT products of the early stages of MT nucleation are extremely heterogeneous (Supplementary Fig. 11), and therefore not suitable for the high-resolution structure determination (Manka and Moores, 2018). Docking of CDC in the conformation found in the crystal structure (Burger et al., 2016) to a minimal (4-dimer) bent tubulin assembly (Supplementary Fig. 10A-C) and the docking refinement by Rosetta illuminates the probability of this hypothesised interaction (Fig. 4B-D and Supplementary Fig. 12). To build the assembly, we used the only available crystal structure of a bent tubulin in the GTP state (PDB: 3RYF) (Nawrotek et al., 2011) and the only available crystal structure of intact CDC (PDB: 5IP4) (Burger et al., 2016), as explained in Supplementary Fig. 10. The lowest-energy complex out of 50 simulated structures shows CDC could form contacts with all 4 tubulin subunits (Fig. 4B-D, Supplementary Fig. 10A-C and Supplementary Fig. 12). We next simulated 50 structures of an NDC complex with the bent tubulin assembly using the same Rosetta protocol (Supplementary Fig. 10B-C), which showed that the best scoring model was worse than the worst scoring models of the equivalent CDC fitting (Supplementary Fig. 10B). We also carried out a control modelling experiment using the experimentally determined NDC complex with lattice-like tubulin as a starting model (as shown in Fig. 2F). The lowest-energy calculated structure also had the lowest RMSD from the experimentally derived NDC-MT structure (Supplementary Fig. 10D), supporting the validity of our modelling approach. All together, these calculations support the idea that CDC, in contrast to NDC, has an intrinsic shape that is compatible with binding to complexes of curved tubulin, and which could predispose it for involvement in initial interactions with nascent MTs (Fig. 2D).

The hypothesised CDC binding to tubulin assemblies would likely accompany the bent-to-straight conformational transition of tubulin (Fig. 4, Supplementary Fig. 12B-D and Movie 1), which is required to convert a pre-MT assembly to MT lattice (Driver et al., 2017; Gigant et al., 2000; Grishchuk et al., 2005; Mandelkow et al., 1991; Nogales et al., 1998; Rice et al., 2008) (Fig. 1A and Supplementary Fig. 11). Such accompaniment of CDC in the vertex of 4 tubulin dimers could help stabilise lateral and longitudinal contacts between these dimers. It might also limit longitudinal flexibility (bending) of the assembly (Fig. 4C and Supplementary Fig. 12B-C), facilitating lattice closure.

The hypothesised and experimental complexes exhibit a logical hierarchy of the interaction area: starting from 758 Å for CDC bound to bent tubulin assembly (modelled), through 1047 Å for CDC bound to GDP.Pi-MT lattice (experimental), to 1603 Å for NDC bound to GDP-MT lattice (experimental) (Fig. 4B). The final CDC and NDC binding to MT lattice likely stabilises tubulin-tubulin contacts and prevents PF curling (and thereby prevents MT depolymerisation) (Fig. 4C). Comparison of the cross-sections of the three complexes: CDC with bent tubulin assembly (modelled) and CDC or NDC with the MT lattice (experimental), allows visual appreciation of the fit between each DC domain and a particular tubulin polymerisation state (Fig. 4D and Supplementary Fig. 12B). It shows how the distinct conformations of loops 1 and 3 in CDC and NDC could mediate distinct interactions with distinct tubulin conformations.

The proposed DCX interaction with the curved pre-MT tubulin oligomers by CDC is also supported by the fact that no MTs were observed when tubulin polymerisation in the presence of NN for 30 seconds was attempted, while a relatively large number of such oligomers were still present after 1 hour-long incubation with NN (Supplementary Fig. 11). Thus, we suggest that MT nucleation by NN is inefficient and architecturally inaccurate compared to WT specifically because of the absence of CDC’s interaction with early tubulin nuclei (Fig. 1D). The high conformational stability of NDC compared to CDC, on the other hand, offers explanation for its apparent role as an MT stabiliser.

### Pathogenic mutations in DC domains affect their fold or interaction with tubulin

Finally, we wanted to investigate how the known symptomatic DCX mutations in human correlate with MT binding interfaces established in this study. We used the OMIM® database (Online Mendelian Inheritance in Man: www.omim.org) to find all pathogenic and likely pathogenic DCX missense mutations, i.e. those that still produce the full length protein, but with point mutations in amino acid sequence that compromise DCX function and cause a disease. We found 78 such mutations of which only 5 do not lie in the DC domains (Supplementary Table 1). The 73 mutations that map to the DC domains involve 65 unique mutation sites, 33 in NDC and 32 in CDC (Fig. 5, red circles). Of the NDC mutation sites, 21 involve residues that contribute to the domain fold (Fig. 5A, white), and 12 are at the MT binding interface (Fig. 5A, green). For the analysis of mutations in CDC, we considered both the experimentally determined MT binding interface (Fig. 2E and Supplementary Fig. 7), and the modelled binding site for a bent tubulin assembly (Supplementary Fig. 12A). Of all CDC mutation sites, 21 involve core residues that determine the domain fold (Fig. 5B, white), 7 are at the MT binding site (Fig. 5B, green with straight numbers) of which 6 overlap with the putative bent tubulin assembly binding site (Fig. 5B, green with straight numbers and dots), and additional 4 are unique to the putative bent tubulin assembly binding site (Fig. 5B, green with *italicised* and shifted numbering). The mapped 21 core residues in CDC are different than the mapped 21 core residues in NDC and they exhibit different distribution within the domain (Fig. 5). Thus, while it is well established that DCX mutations can cause disease by either disrupting DCX binding to MT or by disrupting the DC domain fold (Kim et al., 2003), our data highlight that disruption of domain plasticity may also be a disease mechanism arising from mutations specifically in CDC.

**Figure 5.**
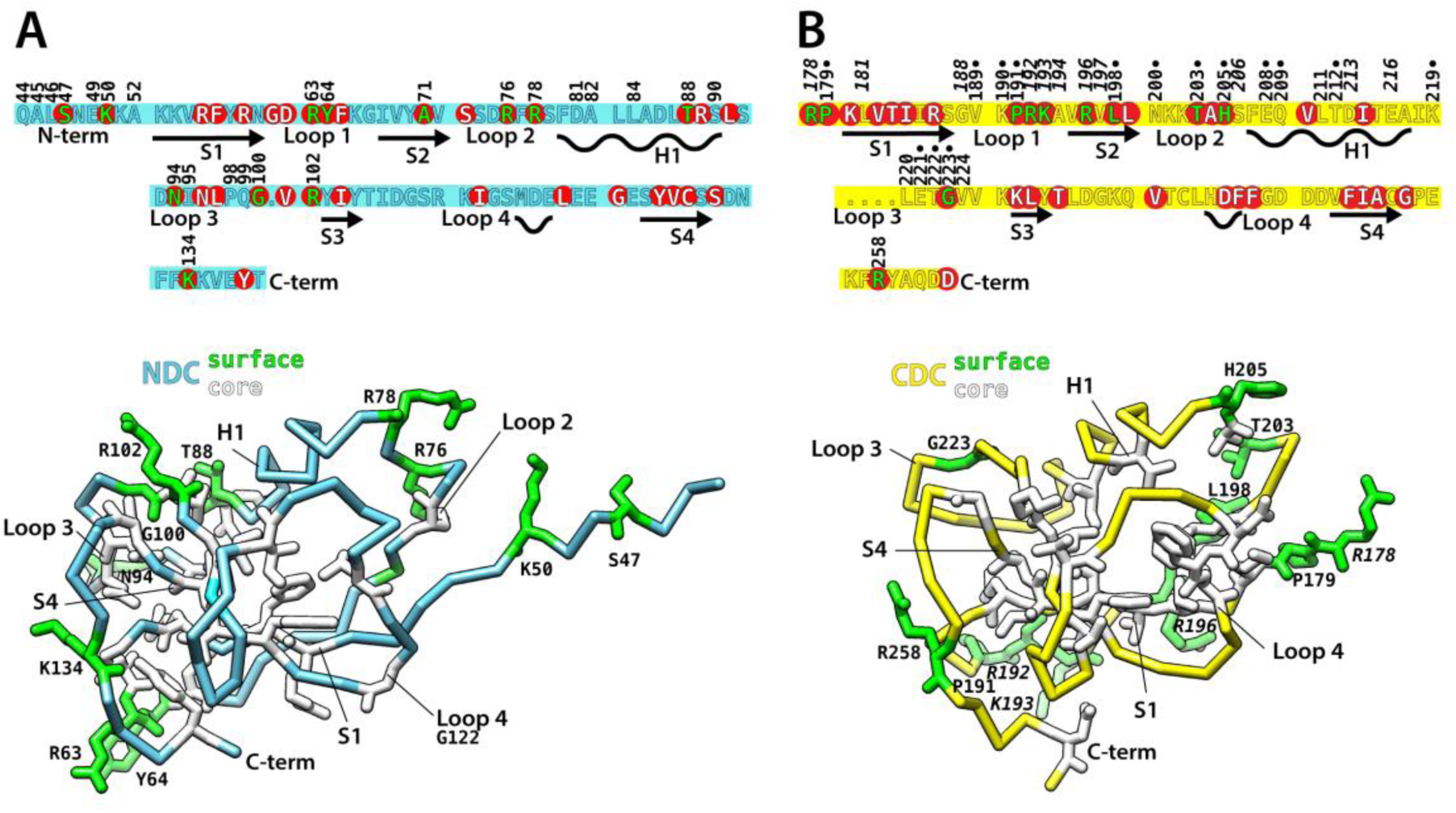
Disease-causing mutations in DC domains map to their core or to DCX-tubulin/MT interface. **A**. The human NDC sequence and the cryo-EM structure of the MT-bound conformation, including the short N-terminal flanking region. Pathogenic mutation sites (Supplementary Table 1) are indicated in the sequence with red circles. Side chains of residues representing the mutation sites are depicted in green if they mediate MT binding or white if they lie in the domain core. MT-binding residues are numbered, and secondary structures are labelled. H, α-helix; S, β-strand. **B**. The human CDC sequence and the cryo-EM structure of the MT-bound conformation. Pathogenic mutation sites (Supplementary Table 1) are indicated in the sequence and the 3D structure as in A. MT-binding residues are numbered using normal font immediately above the sequence and those mediating interactions only with a bent tubulin assembly are indicated with numbering shifted away from the sequence and *italicized*. Residues contributing to both interfaces are indicated with a dot.

## Discussion

DCX is critical for neuronal motility during mammalian brain development. It localises to the distal processes of the migrating immature neurons (Tint et al., 2009), where the absence of the centrosomal γ-tubulin ring complexes (γ-TuRCs) that normally template the 13-PF architecture, may require that DCX contributes to *de novo* MT nucleation to help direct cell migration (Stiess et al., 2010). In this work, we dissect the roles of the two pseudo-repeats (DC domains) in DCX, thereby providing mechanistic insight into how this neuronal migration protein nucleates and stabilises MTs (Fig. 6). DCX needs both DC domains for proper function, but our data suggest that a 2-step mechanism is involved, where each step is served by distinct DC domains. Step 1 is mediated by CDC and pertains primarily to MT nucleation, while step 2 is mediated by NDC as well as CDC and pertains to longer-term MT stabilisation. Crucially, our experiments show that each DC domain can fulfil its entire dedicated role only as a part of the DCX tandem, agreeing with earlier studies (Horesh et al., 1999; Taylor et al., 2000). We observed stimulation by certain single-DC domain constructs on MT growth rates; in addition, and contrary to earlier studies (Bechstedt et al., 2014; Moores et al., 2006), the wild-type DCX (WT), as well as the NDC-NDC tandem (NN), increased MT polymerisation rates at high concentrations, albeit modestly compared to bona fide polymerases such as XMAP215 (Brouhard et al., 2008). The discrepancies between our current work and previous studies likely stem from combinations of the differences between DCX isoforms (Supplementary Fig. 13A) and protein expression systems used in different studies (see further discussions).

**Figure 6.**
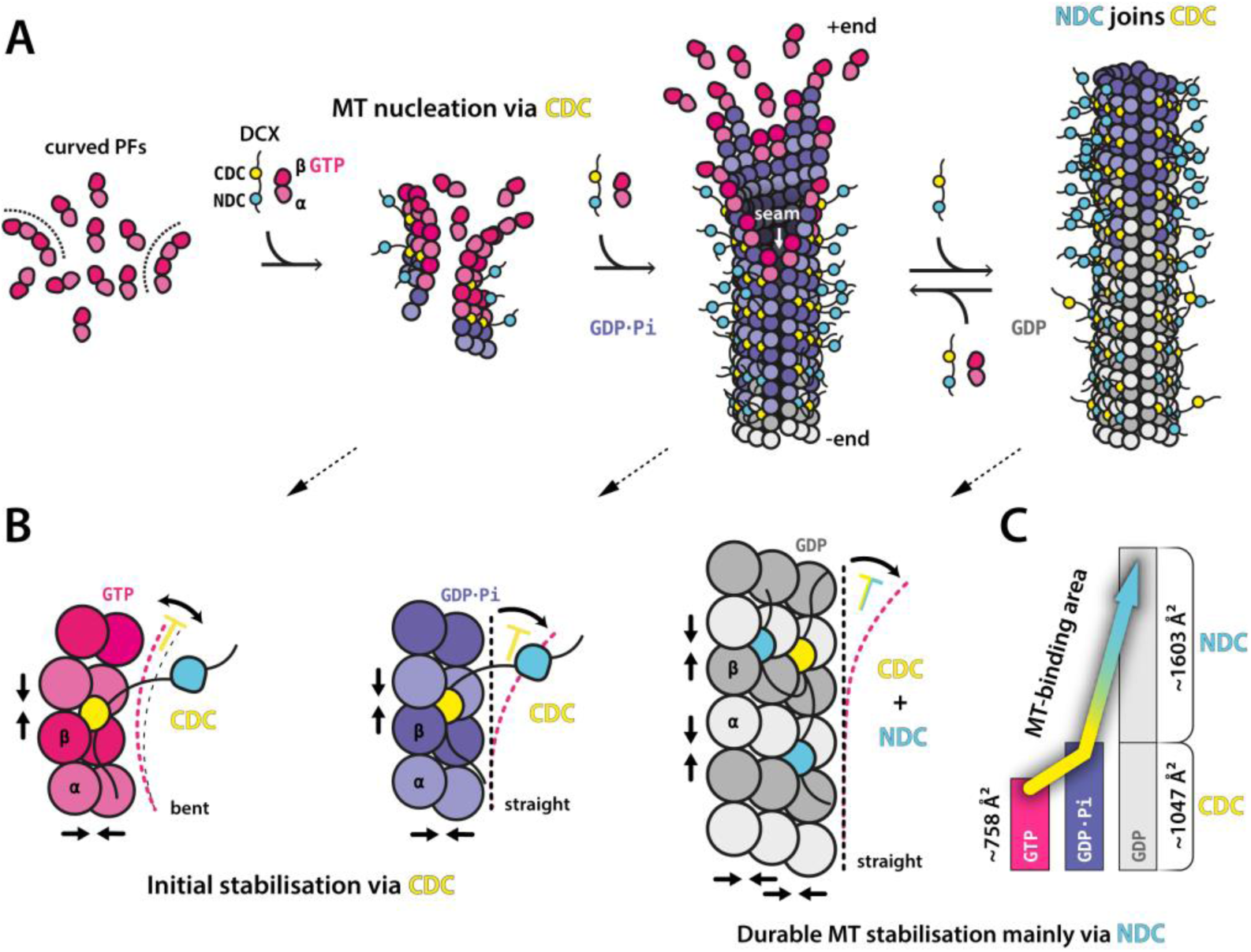
Mechanism of MT nucleation and stabilisation by DCX. **A**. DCX nucleate MTs directly by stabilising early tubulin oligomers (curved PFs). CDC mediates this activity as well as the initial stabilisation of the nascent GDP.Pi-MT lattice. Then NDC joins CDC along the mature GDP-MT lattice. **B**. Summary of the sequential involvement of the two DC domains at different stages of MT assembly, characterised by different (bent or straight) tubulin conformations and different nucleotide states. CDC stabilises tubulin-tubulin contacts (vertical and horizontal arrows) in pre-MT GTP-tubulin oligomers and potentially reduces (T shape) their longitudinal bending (curved arrow), before stabilising tubulin-tubulin contacts in the straight GDP.Pi-MT assembly. CDC appears to remain associated with tubulin assembly during its transition from bent to straight conformation due to its intrinsic structural plasticity. Then NDC joins CDC, stabilising tubulin-tubulin contacts in the straight GDP-MT lattice, preventing (T shape) tubulin bending (curved arrows) and PF peeling (MT breakdown). **C**. Evolution of the DCX-tubulin/MT binding interface at different stages of MT assembly. During the initial CDC interaction with tubulin, the contact site between CDC and tubulin increases by ∼28% (from ∼758 Å^2^ (pink bar) to ∼1047 Å^2^ (violet bar)) during tubulin transition from bent (pre-MT) to straight (MT lattice-like) conformation, accompanied by GTP hydrolysis to GDP.Pi. Then, both CDC and NDC mediate the binding with MT. The contact area between NDC and MT lattice is greater by ∼35% (∼1603 Å^2^; top grey bar) than that of CDC, and together, DC domains occupy ∼2650 Å^2^ area on mature MT lattice (grey bars). Interface areas according to Fig 4C.

We propose that CDC in DCX can directly nucleate MTs, because we found that its substitution for another NDC: i) slows down MT nucleation, and ii) results in a heterogeneous MT population (13-and 14-PF MTs). Moreover, we found by modelling that iii) CDC fits well at the junction of 4 bent tubulin dimers and may not only stabilise the longitudinal and lateral contacts between these dimers (a prerequisite for MT lattice formation), but also limit longitudinal flexing of tubulin oligomers (Fig. 6B). A reduction or restriction in outward bending of laterally associating PFs would be advantageous for subsequent MT lattice closure, which requires tubulin straightening. Our results suggest that CDC remains associated with tubulin oligomers while they undergo the dynamic transition from bent to straight conformation (Movie 1), to become a part of MT lattice. The molten globule properties of CDC were previously characterised using NMR (Kim et al., 2003), while CDC has only recently been crystallised in complex with a stabilising antibody (Burger et al., 2016) or as a domain-swapped dimer (Rufer et al., 2018), also reflecting the domain’s flexible nature. Consistently, our CDC-CDC tandem (CC construct) proved to be unstable at 37 °C. CDC’s role as a quality control agent, tuning MT assembly to produce exclusively 13-PF MTs, may be because it can limit lateral as well as longitudinal conformational variability in tubulin nuclei (Fig. 6B). The C-terminal S/P-rich domain was previously found to also have a regulatory role in determining DCX-mediated MT architecture (Moores et al., 2004), an effect that may be direct or mediated via CDC. Here, we observed MT growth stimulation by CDC-S/P-rich constructs (C1 and C2), but not by an isolated CDC, devoid of the S/P-rich domain (C3 and C4). This further emphasises the apparent regulation of DC domain function by DCX’s disordered C-terminal region, while the instability and low activity of the CDC-NDC chimera points to the importance of the sequence context in which each domain is located. The mechanisms of these effects, especially in the context of DCX isoforms and the presence or absence of post-translational modifications (PTMs), are not understood and will be the focus of future work. Overall, we conclude that the remarkable conformational plasticity of CDC reflects the conformationally dynamic nature of its oligomeric tubulin substrate during MT nucleation.

We identified NDC as the primary stabiliser of the MT lattice after CDC-mediated nucleation. It appears to join CDC along the MT shaft shortly after nucleation, but at large DCX excess over available MT binding sites, NDC appears to replace CDC, suggesting that it has a higher MT lattice binding affinity than CDC. This is consistent with the more extended MT-binding interface of NDC compared to CDC (summarised in Fig. 6C) due to both the co-binding of the short pre-NDC region and a deeper penetration of the lattice with the longer loop 3. Physical linkage with CDC ensures proximity of NDC after MT nucleation, facilitating the DC domain enrichment on MT lattice during MT maturation (Fig. 6A). In theory, DCX binding to MT via both DC domains results in a much stronger interaction between each DCX molecule and MT lattice (Fig. 6C). This would predict that in a certain DCX concentration range, the lower the stoichiometry of DCX on the MT lattice, the tighter its affinity. Conversely, the cooperative DCX binding to MT, described by the Brouhard lab (Bechstedt and Brouhard, 2012), means that DCX dissociates from MT lattice more slowly as the DCX concentration increases. This may mean that the cooperative DCX binding mechanism may not be mediated by the DCX molecule itself, but rather via MT lattice effects exerted by DCX, such as the lattice regularisation that may optimise DC domain binding sites.

Unlike CDC, NDC has been crystallised multiple times (Burger et al., 2016; Cierpicki et al., 2006; Kim et al., 2003), and superposition of these X-ray structures with our cryo-EM structure suggests high conformational stability of NDC, which suits its proposed role as an MT anti-catastrophe agent. Such rigidity combined with the precise fit to MT lattice vertex prevents PFs from curling outward and breaking apart. Each such vertex is built from four tubulin dimers, and – in saturating conditions - each tubulin dimer in the lattice can be held by four NDC domains. Despite being relatively conformationally rigid, we found NDC still capable of adapting to slightly varying inter-PF angle, as evidenced with equal decoration of 13- and 14-PF MTs with NDC-NDC tandem (NN construct). This property of NDC – enabled primarily through loop flexibility - has not previously been reported, although the first low-resolution cryo-EM reconstruction of MT decorated with C-terminally truncated DCX (tDCX, lacking the S/P-rich region) clearly showed a DC domain binding to the 14-PF lattice (Moores et al., 2004). The potential physiological significance of this adaptation is not known.

Our sequential mechanism of MT nucleation and stabilisation by DCX is overall consistent with earlier work (Bahi-Buisson et al., 2013; Bechstedt and Brouhard, 2012; Bechstedt et al., 2014; Ettinger et al., 2016; Fourniol et al., 2010; Horesh et al., 1999; Liu et al., 2012; Moores et al., 2004, 2006; Sapir et al., 2000; Taylor et al., 2000). Previous low- and intermediate-resolution cryo-EM studies proposed NDC as the predominant MT-binding domain (Fourniol et al., 2010; Liu et al., 2012; Moores et al., 2004). Using 366 amino-acid (aa) isoform of human DCX (a C-terminal splice variant (Supplementary Fig. 13A) expressed in eukaryotic cells, the Moores lab visualised DCX density bound to MTs that resembled NDC (including the pre-NDC flanking region) in the presence of a kinesin motor domain (Fourniol et al., 2010; Liu et al., 2012). However, decoration with DCX alone showed no density for the pre-NDC flanking region but did reveal docking of the post-NDC linker region along the NDC core (Liu et al., 2012), likely via W146 as shown earlier in solution (Cierpicki et al., 2006). In our current study using bacterially expressed 360 aa human isoform 2 (Supplementary Fig. 13A), we did not observe such linker docking, which may be due to the isoform differences and/or the lack of PTMs that may play additional regulatory roles.

A co-pelleting study using MTs pre-stabilised by Taxol reported that only a CDC-specific, and not an NDC-specific, antibody prevented DCX from binding MTs, suggesting that it is primarily CDC that mediates the DCX-MT interaction (Burger et al., 2016). However, we previously found that Taxol MT stabilisation reduces subsequent DCX binding (Manka and Moores, 2018), while others have found that addition of Taxol strips DCX (expressed in physiological levels) from MTs in cells, except from curved MT regions (Ettinger et al., 2016). Our new data suggest that Taxol MTs may support CDC - but not NDC - binding to MTs through its recognition of bent or flexible MT lattice regions, an interaction that is inhibited by anti-CDC antibodies. The apparent inhibition of NDC binding to straight MT lattice by Taxol is enigmatic, but - as we previously suggested (Manka and Moores, 2018) - may be related to the drug’s mechanism of MT stabilisation, producing a looser MT lattice and more flexible MTs (Kellogg et al., 2017; Kikumoto et al., 2006).

Our analysis of the disease-linked missense mutations in the DC domains reveals that these mutations affect either the domain fold or the hereby determined DCX-MT interaction. Interestingly, the 65 sites in the DC domains identified as essential for DCX function appear to be completely conserved across the vertebrate classes, in which forebrain is the dominant part of the brain: from mammals, through birds and reptiles, to amphibians (Supplementary Fig. 13B-D). We found divergence in 6 of these residues in fish, the lowest vertebrate, where the midbrain is the dominant part of the brain: 3 in NDC and 3 in CDC (Supplementary Fig. 13C-D). This suggests that the observed adjustments in the sequence of the DC domains may have a role in enabling forebrain development in higher vertebrates, potentially contributing to wider molecular synergies in this part of the brain and elaboration of the mammalian neocortex (Briscoe and Ragsdale, 2018).

The characterisation of distinct roles for each DC domain within DCX reflects the conformational diversity of tubulin as it transitions between unpolymerised and polymerised states (Brouhard and Rice, 2018), and is influenced by MT aging, damage and repair (Schaedel et al., 2015). In this context, different domains within MAPs effectively act as stage-specific tubulin chaperones specialising in interactions with different tubulin conformations. The presence of DC domain pseudo-repeats with different functional specialisations supports multiple activities in DCX, including acceleration of MT nucleation, control of MT architecture, MT stabilisation, and interaction with other components of MT cytoskeleton (Liu et al., 2012), all within a single molecule. It is also possible that DCX facilitates MT repair by chaperoning GTP-tubulin oligomers destined for incorporation into the site of MT damage via CDC and by subsequently stabilising the repair site via NDC.

The presence of pseudo-repeats is common in MAPs, for example the TOG domains in XMAP215/CLASPs (Cook et al., 2019; Slep, 2018), STOP or MAP2/tau family of MAPs containing 3-4 pseudo-repeats (Feng and Walsh, 2001). This reflects the importance for many of these proteins of (transiently) sensing or stabilising a conformational range of tubulin ensembles undergoing dynamic transitions. Such modular MAPs can also favour or disfavour interactions with other MAPs, motors (kinesin and dynein) or MT modifying enzymes. They also regulate MT bundling or cross-linking with other cytoskeletal assemblies such as actin filaments. It will be interesting to evaluate the differential roles of pseudo-repeat containing binding proteins in other cytoskeleton filament systems in the light of our findings.

## Methods

### Generation and cloning of DCX variants

Doublecortin (DCX) constructs were based on human isoform 2 sequence (Uniprot identifier: 043602-2) – here called wild-type (WT) – and designed through inspection of the available crystal structures. The truncated variants were generated using standard PCR methods from the gene sequence kindly gifted by Anne Houdusse (Institute Curie, Paris). Constructs were subcloned into pNic28-Bsa4 vector (Structural Genomics Consortium, Oxford, UK), with an N-terminal tobacco etch virus (TEV) protease-cleavable His-tag. The DC domain duplication and swapped chimeras were obtained with a multistep PCR amplifications and restriction enzyme reactions (New England Biolabs) as depicted in Supplementary Fig. 1.

### Purification of DCX variants

All DCX constructs were expressed in BL21 Star (DE3) *E. coli* cells (Invitrogen) at 18 °C. The cells were spun and resuspended in ice-cold lysis buffer (50 mM Na2HPO4 pH 7.2, 300 mM NaCl, 10 mM imidazole, 10% glycerol, 2 mM DTT) supplemented with protease inhibitor cocktail (cOmplete Cocktail Tablet, Roche/Sigma Aldrich). The cells were lysed by sonication. The lysates were clarified by centrifugation and passed through a nickel HisTrap HP column (GE Healthcare). The His-tagged proteins were then eluted with 10-250 mM imidazole gradient. To cleave off the tags we used a His-tagged TEV protease expressed in-house and both the cleaved His-tag and the His-tagged protease were removed from protein solutions by passage over loose Ni-NTA His-bind resin (Merck). The tag-free DCX variants were then captured on a HiTrap SP HP ion exchange column (GE Healthcare) equilibrated in BRB80 buffer (80 mM PIPES [piperazine-N,N′-bis(2-ethanesulfonic acid)] pH 6.8, 1 mM EGTA [ethylene glycol-bis(β-aminoethyl ether)-N,N,N’,N’-tetraacetic acid], 1 mM MgCl2, 1 mM DTT [dithiotreitol]) and eluted with NaCl gradient (15-300 mM). Final purification and desalting were done by gel filtration through Superdex 200 size exclusion column (GE Healthcare) equilibrated in BRB80 buffer.

### Thermal shift protein stability assay

The stability of DCX constructs was assessed with temperature denaturation monitored by SYPRO Orange dye that becomes highly fluorescent on contact with exposed hydrophobic regions (ThermoFluor). The 5000X SYPRO Orange dye solution in DMSO (Life Technologies) was diluted 250 times in BRB80 buffer containing 5 μM DCX construct. The change in fluorescence was measured in a 96-well plate by MyIQ RT-PCR instrument (BioRad). The melting temperature (T_m_) is at the inflection point of the upward curve and is conveniently identified with a peak of a derivative function (Supplementary Fig. 3B-C).

### Turbidimetric MT nucleation assay

MT nucleation can be monitored by the extent of light scattering or turbidity. Tubulin at 5 μM concentration was mixed with 1-5 μM DCX construct in BRB80 buffer in an Eppendorf UVette (220-1600 nm). The increasing sample turbidity was measured at 500 nm wavelength over 30 min at 37 °C with Cary 3 UV/VIS spectrophotometer equipped with a temperature control unit (Varian).

### Negative-stain EM

Samples removed directly from turbidimetric assays were deposited on EM grids with a continuous carbon film (Agar). The grids were briefly blotted and washed with BRB80 buffer before staining with 1% uranyl acetate solution in water (negative stain solution). After ∼1 sec exposure to uranyl acetate the grids were blotted and air dried. The negatively-stained grids were imaged on a 120 kV Tecnai T12 microscope (FEI) with a US4000 4K x 4K CCD camera (Gatan).

### TIRF microscopy assays

All TIRF microscopy assays were performed in ∼10 μl flow chambers built through attachment of biotin-PEG coverslips (Stratech) to glass slides with a double-sided tape. The chambers were blocked by 5 mg/ml casein solution in 0.75% Pluronic F-127, washed with 0.4 mg/ml casein solution in BRB80 buffer (Wash solution), coated with 0.5 mg/ml neutravidin solution in BRB80 buffer, and washed again with Wash solution before application of GMPCPP-MT seeds. The fluorescent and surface-attaching (neutravidin-binding) GMPCPP-MT seeds were polymerized from a mixture of 10 μM unlabelled tubulin, 10 μM Alexa488-tubulin, 10 μM biotin-tubulin (Cytoskeleton) and 1 mM GMPCPP (Jena Biosciences) in BRB80 buffer, incubated at 37 °C for 30 min.

For MT stabilization assay, ∼0.1 μM GMPCPP-MT seed solutions in BRB80 (according to tubulin dimer concentration) were applied to flow chambers. After washing with Wash solution the chambers were ready to receive an assay solution comprising: 9 μM unlabelled tubulin, 1 μM Alexa488-labelled tubulin, 0.4 mg/ml casein, 1 mM GTP, 5 mM DTT, 20 mM glucose, 0.1 % methyl cellulose, 300 μg/ml glucose oxidase and 60 μg/ml catalase, with varying concentrations of unlabelled DCX variant in BRB80 buffer.

MT dynamics was observed using an Eclipse Ti-E inverted microscope equipped with a CFI Apo TIRF 1.49 N.A. oil objective, Perfect Focus System, H-TIRF module, LU-N4 laser unit (Nikon) and a quad band filter set (Chroma). Exposures were recorded over 100 ms with 2 sec intervals or continuously on an iXon DU888 Ultra EMCCD camera (Andor), using NIS-Elements Software (Nikon). Kymographs of dynamic and DCX-stabilized MTs were generated in FiJi (https://fiji.sc/>).

### Cryo-EM sample preparation

Tubulin stock was prepared by reconstituting lyophilised bovine brain tubulin (Cytoskeleton) to 100 μM concentration in BRB80 buffer supplemented with 1 mM GTP.

For GDP.Pi-MTs 10 μM GTP-tubulin was mixed with 50 μM WT in cold BRB80 buffer containing 1 mM GTP. The mixture was immediately applied to a glow-discharged Lacey grid (Agar) and transferred to a Vitrobot (FEI/Thermo Fisher Scientific) for rapid on-grid MT polymerisation in a humid and warm (30 °C) Vitrobot chamber. After 30 sec incubation, the grid was blotted and plunge-frozen in liquid ethane.

For GDP-MTs 5 μM GTP-tubulin was co-polymerized for 30 min at 37 °C with 3 μM WT or NN in BRB80 buffer containing 1 mM GTP. These MTs were applied to glow-discharged Lacey grids and incubated for 20 sec at room temperature. Then the grids were briefly blotted and 50 μM WT or NN solution in BRB80 buffer was added to maximize MT decoration. The grids were then transferred to the Vitrobot and incubated there for 1 min at 30 °C, before being blotted and vitrified as before.

### Cryo-EM data collection

Cryo-micrographs were acquired on a 300 kV Polara microscope (FEI) with a K2 Summit camera (Gatan) operated in counting mode after a Quantum energy filter (Gatan) with a 20 eV slit. The magnified pixel size was 1.39 Å. The dose rate was 2.6-2.8 e-/Å^2^/sec during 9 sec exposures, resulting in the total dose of 23-25 e-/Å^2^ on the specimen. These exposures were collected manually and fractionated into 36 movie frames (0.25 sec/frame) with SerialEM (http://bio3d.colorado.edu/SerialEM/), at defocus ranging from −0.4 to −2.5 μm defocus.

### Cryo-EM image processing and 3D reconstruction

Movie frames were aligned using MotionCor2 (Zheng et al., 2017), with images divided to 25 tiles for local correction of beam-induced motion. Start and end positions for each uninterrupted MT were manually selected using the helix mode in EMAN 1 Boxer (Ludtke et al., 1999) that automatically generated 652 x 652 pixel boxes with 474 pixel overlap along MTs. This box size corresponded to MT segments spanning ∼11 tubulin dimers with ∼8 dimer overlap between the segments.

For 3D reconstruction of 13-PF MTs, the MT segments boxed with EMAN1 were averaged in Spider (Frank et al., 1996). Segment averages revealing: i) MT architecture other than 13-PF (in the case of NN-MTs), ii) lattice defects and/or iii) poor contrast (e.g. blurring) were excluded from further processing. The selected high-quality 13-PF segments were treated as single particles in Chuff (Sindelar and Downing, 2007), a custom-designed multi-script processing pipeline using Spider (Frank et al., 1996) and Frealign (Grigorieff, 2007). The initial 2D alignment was aimed at identifying the MT seam location by projection matching in Spider. The reference projections for this step were generated using a synthetic 13-PF DCX-MT reference volume filtered to 30 Å. The contrast transfer function (CTF) parameters for each micrograph were estimated with CTFFIND3 (Mindell and Grigorieff, 2003), and the CTF correction was performed during local refinement within Frealign, producing isotropic 3D reconstructions with pseudo-helical symmetry applied 12 times. This symmetry operation was chosen to average only the PFs occupied with DCX, excluding the unoccupied seam. Independently processed half maps were combined and sharpened in Relion 1.4 (Scheres, 2012) through its automated post-processing routine. The average resolutions of the final maps were estimated using 0.143 FSC criterion and the absence of over-fitting was confirmed with high-resolution noise substitution test (Chen et al., 2013) (Table 1, FSC_true_).

For 3D classification and refinement of 13- and 14-PF NN-MTs, the MT start and end coordinates selected using EMAN 1 (as above) were imported to Relion 2.1 (Kimanius et al., 2016), and 432 x 432 pixel MT segments spaced with 1 dimer distance were generated. 3D classification was done with 15 Å^2^ resolution cut-off using 4x binned segments (resulting in 5.56 Å^2^ pixel size) against 6 synthetic volumes representing 11-, 12-, 13-, 14-, 15-, and 16-PF MT architectures. The most populated 13-PF and 14-PF classes (together containing 99% of all MT segments) were then separately refined, applying pseudo-helical symmetry 12 times (for 13-PF class) or 13 times (for 14-PF class). The resultant 3D maps were sharpened according to local resolution and the average resolution was estimated as before.

### Determination of relative NDC occupancy on different MT architectures

Fourier transforms of the relevant MT segments were averaged using EMAN 1 (Ludtke et al., 1999). Intensities of 8 nm layer lines corresponding to MT-binding periodicity of NDC in relation to 4 nm layer lines corresponding to tubulin subunit periodicity were then calculated using FiJi (https://fiji.sc/). To estimate the error of these ratios we divided each dataset into 3 approximately equal subsets and calculated the average ratio and standard deviation within each set of 3 subsets.

### Atomic model refinement

The 1MJD NMR structure of human NDC (Kim et al., 2003) or the 5IP4 crystal structure of human CDC (Burger et al., 2016) were combined with previously refined tubulin subunits (Manka and Moores, 2018) to create starting models for MT-bound DC domain refinements in our cryo-EM maps. The maps were zoned in UCSF Chimera (Pettersen et al., 2004) around DC domains surrounded by 4 (2 β and 2 α) tubulin subunits. Each isolated map was placed in a new unit cell with P1 space group. The model-map fit was adjusted in Coot (Emsley et al., 2010) before 10 macro cycles of refinements in real space were carried out using phenix.real_space_refine (http://phenix-online.org/) (Afonine et al., 2018) with default settings. Phenix automatically determined weight between data and the restraints to achieve RMS deviations of covalent bonds not greater than 0.01, and for angles not greater than 1.0 from ideal values. Manual adjustments of poorly fitting regions in Coot followed by real space refinements described above were repeated until a satisfactory level of model:map agreement and excellent model geometry were accomplished (Table 1).

### Rosetta-assisted modelling

Two longitudinally associated GTP-tubulin dimers in a bent conformation were extracted from the 3RYF crystal structure (Nawrotek et al., 2011). Two copies of this dimer of tubulin dimers were superimposed on two laterally connected DCX-binding β-tubulin subunits derived from our cryo-EM structure of the 13-PF DCX-MT, as illustrated in Supplementary Fig. 10A. To decrease the global conformational search space and improve the efficiency of the subsequent local docking process in Rosetta (Gray et al., 2003), the 5IP4 crystal structure of human CDC was manually positioned in the vertex of the 4 assembled tubulin dimers in an orientation similar to that observed for CDC bound to GDP.Pi-MT lattice, within 2-6 Å distance from each tubulin subunit. We first ran the Rosetta *relax* protocol, to relieve clashes in the assembly and to move the assembly to the nearest local minimum in the energy function. We then generated 50 model complexes applying the full Rosetta local docking protocol with fixed backbones, specifying CDC as the docking partner for the bent tubulin assembly. The protocol involves course centroid movements, all-atom refinements with side chain packing and random Monte Carlo translations and rotations (Gray et al., 2003). Each individual simulation started with random coarse perturbation of CDC by 3 Å translation and 8° rotation. At the refinement step translation and rotation magnitudes were limited to 0.1 Å and 5°, respectively. Side chains from the input coordinates (pdb) files were added to the rotamer libraries. Each simulation ended with minimization of the interface. The same protocol was employed to generate 50 models of NDC docked to the same bent tubulin assembly and 50 control models of NDC docked to MT lattice-like tubulin assembly, starting from the cryo-EM structure of MT-bound NDC. Each experiment resulted in clear convergence between RMSD and the total energy score (docking funnel), which indicates the validity of the results.

### Structure analyses and presentation

Analyses and visualisations of our cryo-EM density maps and the atomic models refined in those maps were done using PDBePISA (http://www.ebi.ac.uk/msd-srv/prot_int/cgi-bin/piserver), PyMOL (Schrödinger), UCSF Chimera (Pettersen et al., 2004) and ChimeraX (Goddard et al., 2018).

## Data availability

Our maps and coordinates were deposited in EMDB/PDB with the following accession codes: 13-PF WT(CDC)-GDP.Pi-MT (EMD-4861; PDB 6RF2), 13-PF WT(NDC)-GDP-MT (EMD-4858; PDB 6REV), 13-PF NN(NDC)-GDP-MT (EMD-4862; PDB 6RF8), 14-PF NN(NDC)-GDP-MT (EMD-4863; PDB 6RFD).

## Acknowledgements

This work was funded by grants from the Medical Research Council, U.K. to C.A.M (MR/R000352/1). EM data collection was supported by grants from the Wellcome Trust (079605/Z/06/Z, 101488/Z/13/Z) and the BBSRC (BB/L014211/1). Anthony Roberts provided essential advice concerning TIRF microscopy while Joe Atherton and Alex Cook provided valuable guidance about MT processing using RELION.

## Author Contributions

S.W.M. conceived experimental strategies, designed and carried out experiments and computations, analysed data, interpreted results and wrote the manuscript; C.A.M. proposed and supervised the research, interpreted results and wrote the manuscript.

## Supplementary Information

**Supplementary Fig. 1.**
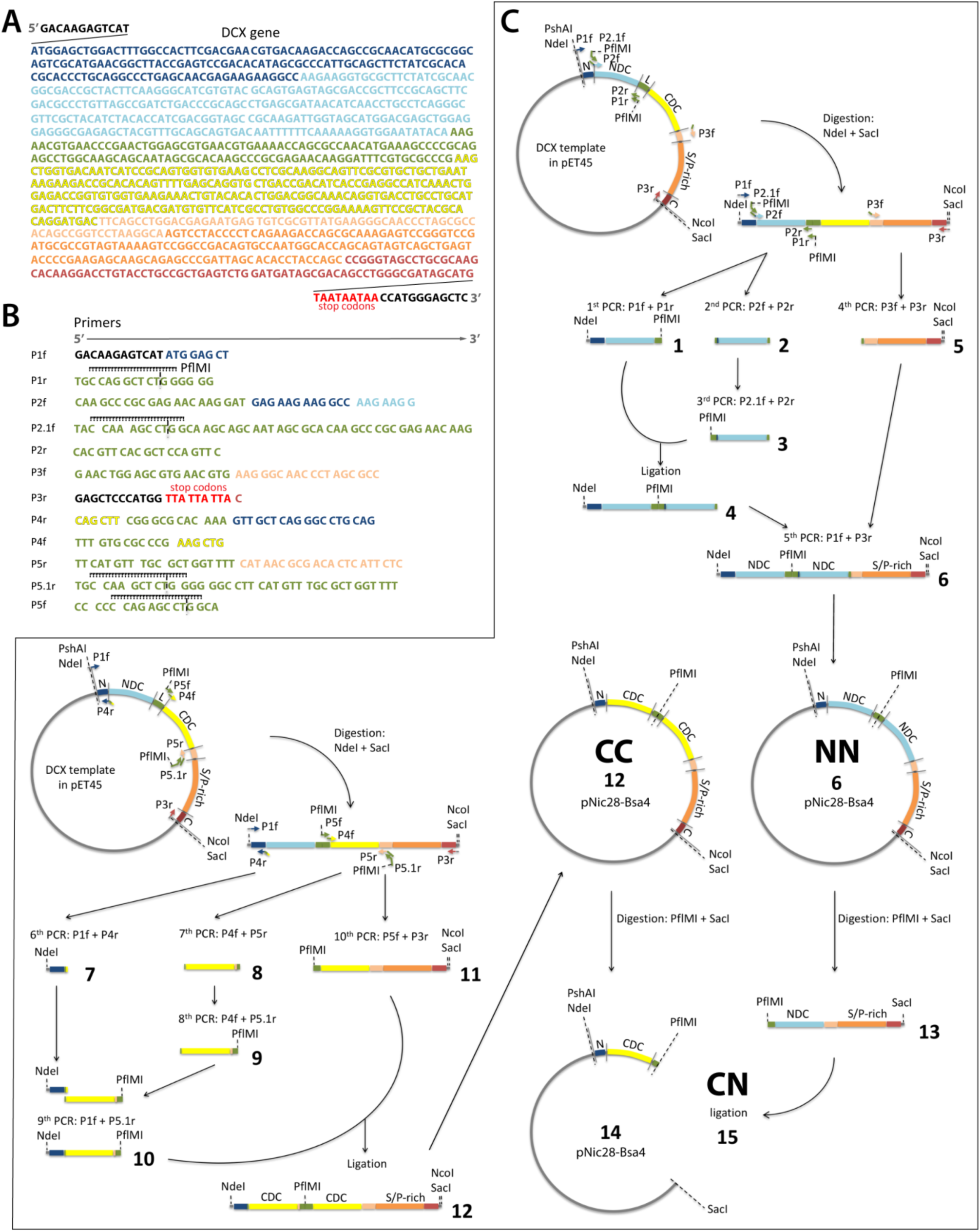
Cloning of the DCX constructs with repeated or swapped DC domains. **A**. Sequence of DCX isoform 2 gene used in this study with flanking sequences covering primer binding sites. Coloured according to region: navy blue, N-terminal region; blue, NDC; green, linker; yellow, CDC; light orange, post CDC region; orange, S/P-rich domain; dark red, C-terminal region. **B**. List of primers used in the generation of the chimeras. Recognition and cleavage site for PflMI endonuclease is indicated. **C**. Schematic representation of the PCR, restriction enzyme digestion and ligation steps (1-15) in the generation of the chimeras. NDC-NDC chimera, NN; CDC-CDC chimera, CC; CDC-NDC chimera, CN.

**Supplementary Fig. 2.**
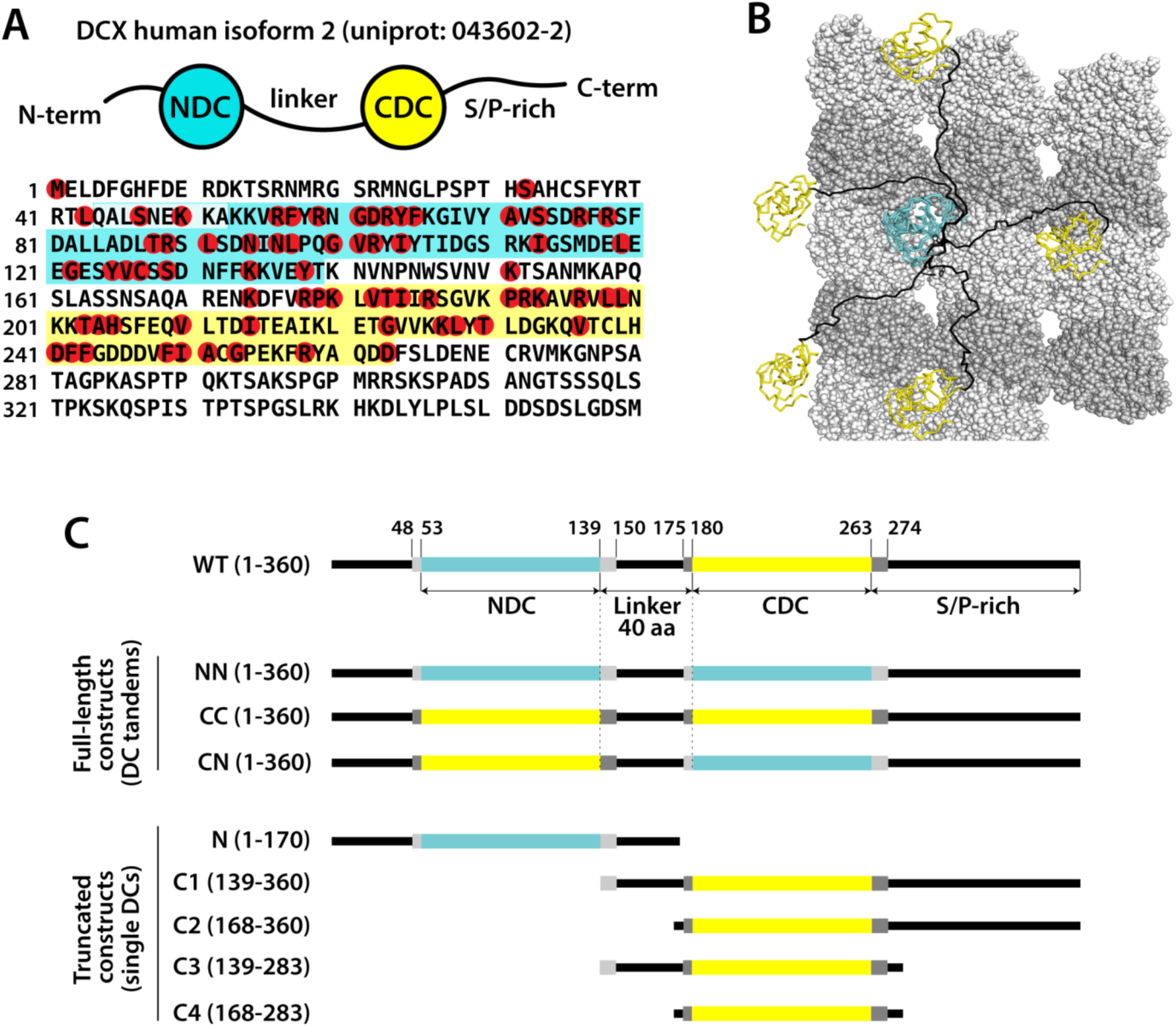
Characteristics of the DCX structure and details about the generated constructs. **A**. Schematic of DCX structural regions and the location of symptomatic missense mutation sites according to the OMIM.org database**. B**. Manual modelling (*sculpting*) in PyMOL shows that the length of the DCX linker region (black) is in principle compatible with multiple modes of DCX binding to MT lattice. NDC, blue; CDC, yellow; *α*-tubulin, dark grey; *β*-tubulin, light grey. **C**. List of DCX constructs generated in this study with database (Uniport.org) DC domain boundaries indicated with colour and the cloning boundaries (including flanking regions) indicated with grey zones. We took care to maintain the length of the linker (dotted lines) in all full-length constructs (DC tandems).

**Supplementary Fig. 3.**
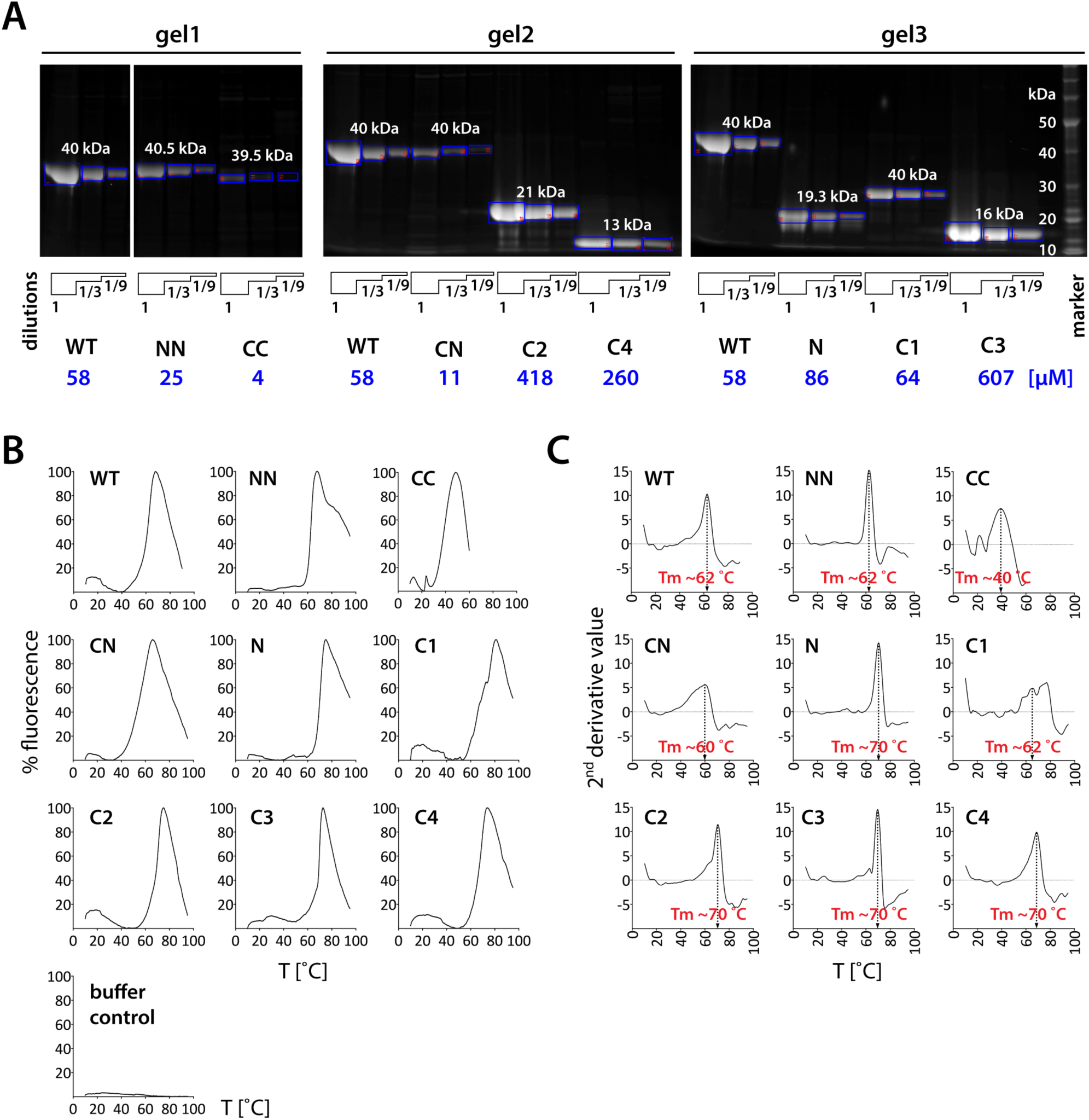
Purity and thermal stability of the generated constructs. **A**. Purity of DCX proteins was assessed by SDS-PAGE under reducing conditions using Bolt Bis-Tris Plus gels (Thermo Fisher Scientific). Peak fractions from the S200 gel filtration column (final purification step) were pooled, concentrated using Vivaspin columns (GE Healthcare) and run neat and at two indicated dilutions (1, 1/3 and 1/9). All gels included WT sample as an internal standard. Some lanes in gel 1 were removed as irrelevant to this study. The gels were stained with SYPRO Orange Protein Gel Stain (Sigma-Aldrich) and scanned using Fujifilm FLA-3000 Fluorescence Laser Imaging Scanner. The intensities of the bands were quantified using FiJi (https://fiji.sc/; as indicated with the blue boxes with red annotations), multiplied according to their dilution factor and averaged together. Protein concentrations are derived from the band intensity ratio to the WT internal standard, which was measured by absorbance at 280 nm wavelength. **B**. Normalized thermal shift (ThermoFluor) plots. The inflection point on the rising curve is the melting temperature (T_m_)**. C**. Second derivative plots to identify T_m_ with the peaks, corresponding to the inflection points in the thermal shift curves in D.

**Supplementary Fig. 4.**
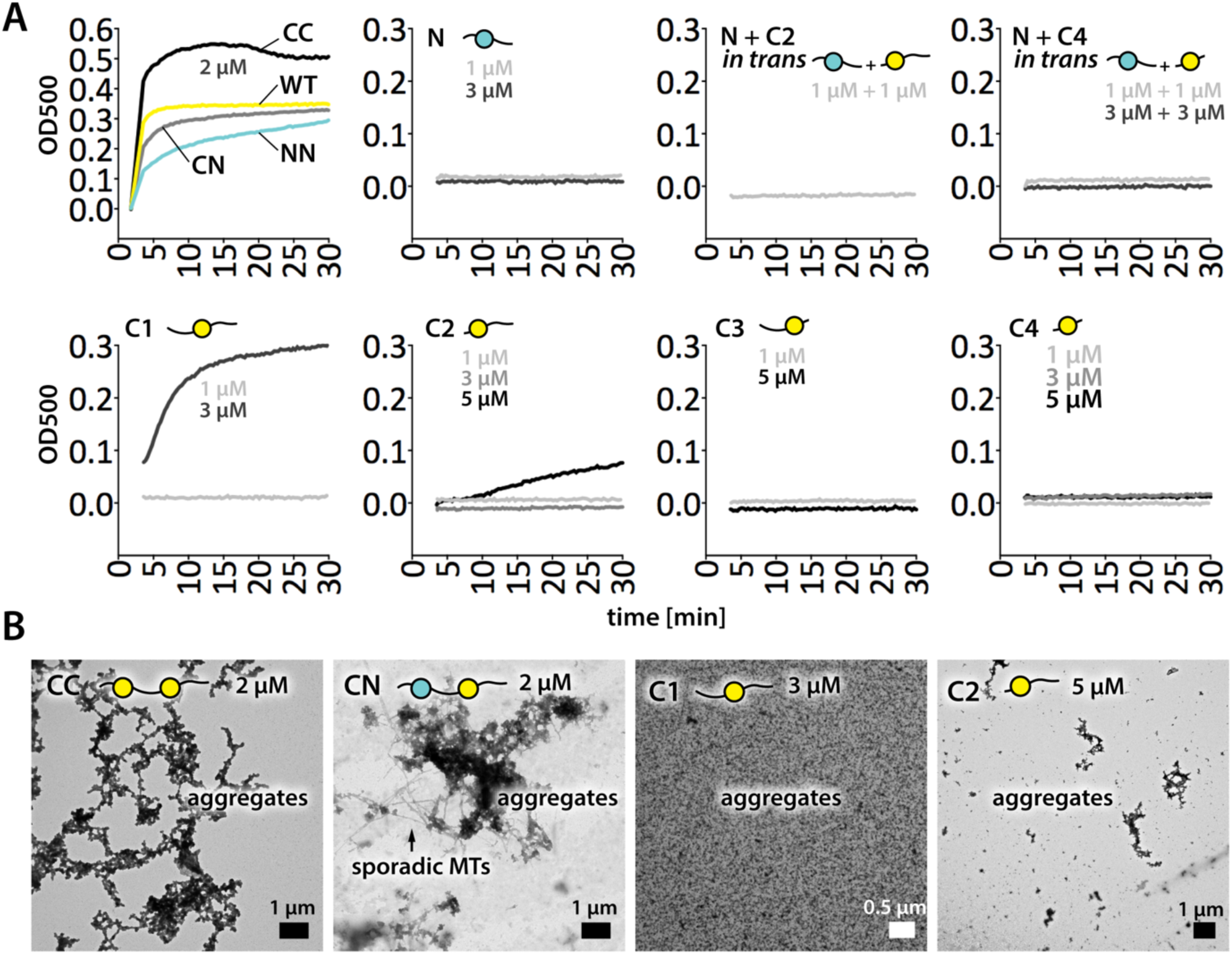
MT nucleation, abilities of different DCX constructs. **A**. Turbidity assay plots. 5 μM tubulin was mixed with indicated concentrations of various DCX constructs and light scattering was measured at 500 nm wavelength (OD500). **B**. Direct verification of turbidity assay results by negative stain EM imaging of selected samples from A. Protein aggregation increases sample turbidity and can be confused for MT nucleation in this assay.

**Supplementary Fig. 5.**
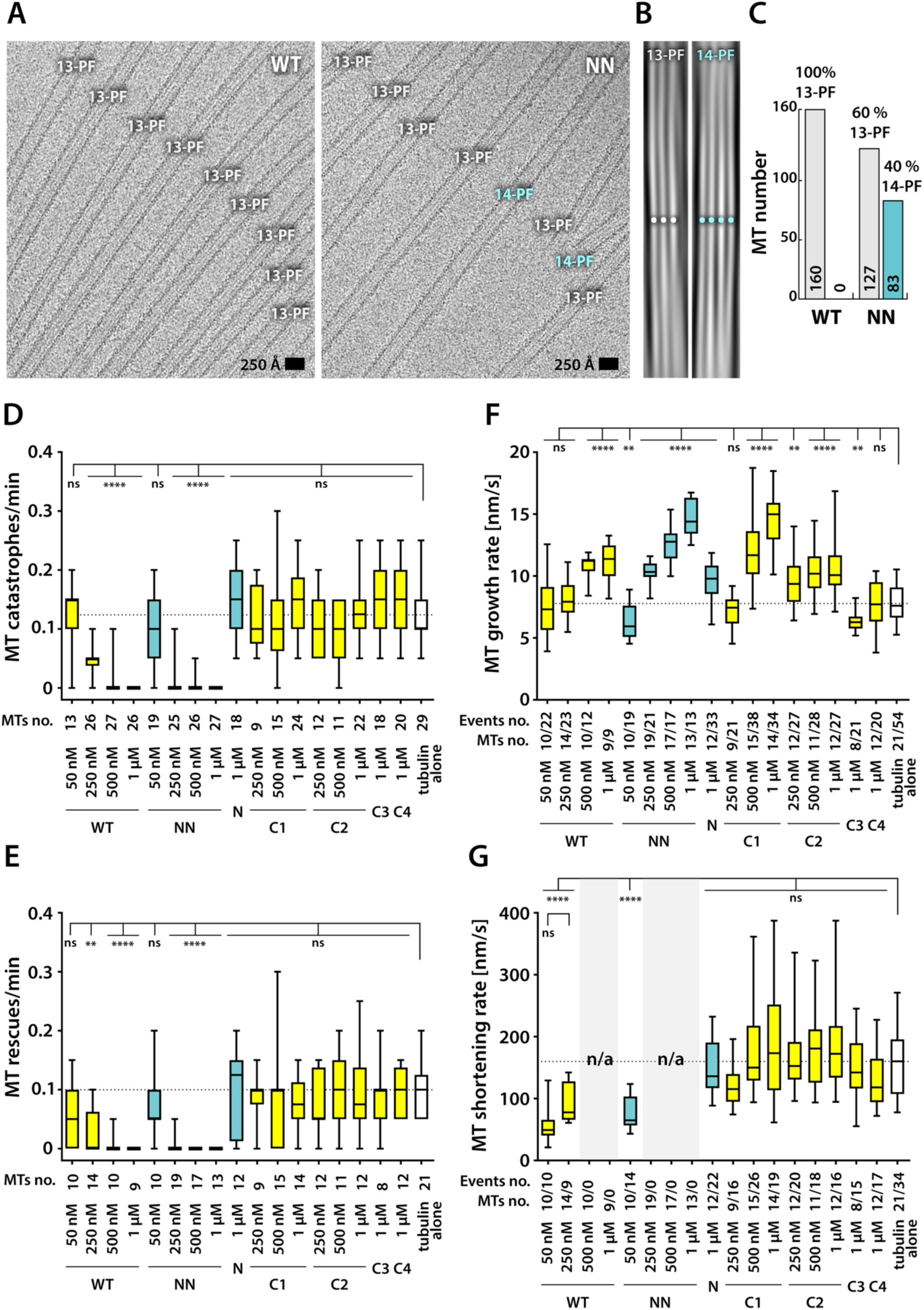
MT populations stabilised by WT and NN and the influence of all DCX constructs on MT dynamic instability parameters. **A**. Example cryo-EM micrographs of WT-MTs and NN-MTs presenting moiré patterns corresponding to different (13-PF or 14-PF) MT architectures. 14-PF MTs are only found in the NN-MT sample. **B**. Example Fourier filtering in FiJi (fiji.sc), depicting differences in moiré patterns of 13-PF and 14-PF MTs, used as a diagnostic in this study. **C**. Quantification of PF number distribution in WT-MT and NN-MT samples. **D-G**. Box & whiskers plots presenting the influence of different DCX constructs on MT dynamic instability parameters measured with TIRF microscopy. The whiskers are defined by the minimum and the maximum measured value, the box shows the interquartile range (between the 25^th^ and 75^th^ percentile) and the central line indicates the median value (50^th^ percentile). The sample sizes: number of MTs and, where appropriate, events analysed are indicated. Statistical analysis was done using ordinary one-way ANOVA and significance has been verified through Holm-Sidak’s multiple comparisons test within Prism 8 (graphpad.com). Stars indicate significant differences compared to the reference value of tubulin alone according to the multiplicity adjusted p values: ****, p < 0.0001; ***, p < 0.0005; **, p < 0.01; *, p < 0.05; ns, not significant (p > 0.05). Tubulin concentration was 10 μM in all experiments.

**Supplementary Fig. 6.**
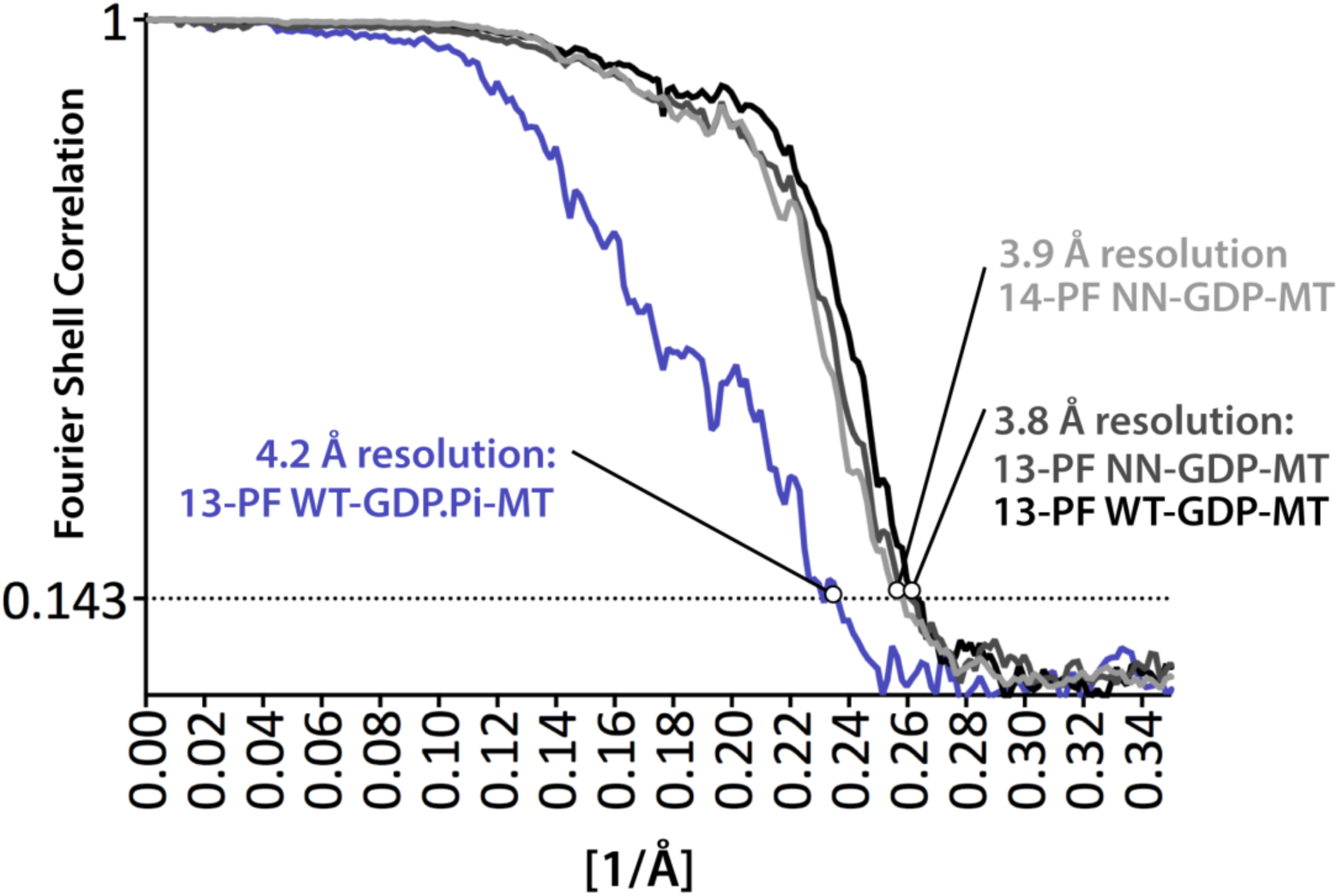
Resolution estimation of cryo-EM reconstructions. Fourier Shell Correlation (FSC) plots and resolution estimation by the 0.143 cut-off criterion.

**Supplementary Fig. 7.**
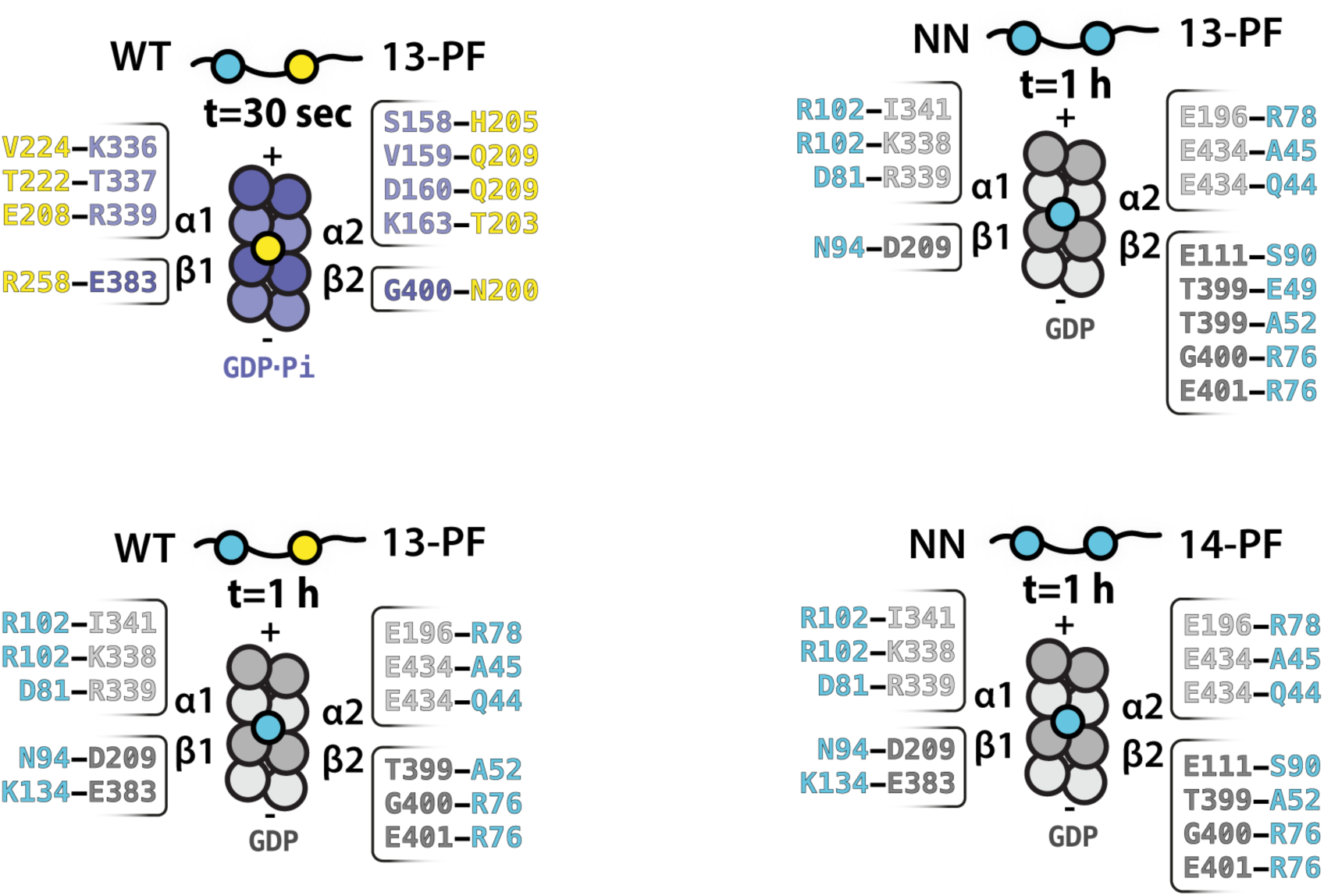
Summary of the resolved WT-MT and NN-MT polar contacts. Schematic diagrams of DC domains bound in the vertices of 4 tubulin dimers in WT-MTs and NN-MTs, according to our cryo-EM reconstructions, including listing of polar contacts (H-bonds and/or salt bridges) made by these DC domains with indicated tubulin subunits. The contacting residues were identified with help of PDBePISA (http://www.ebi.ac.uk/msd-srv/prot_int/cgi-bin/piserver). NDC, blue; CDC, yellow.

**Supplementary Fig. 8.**
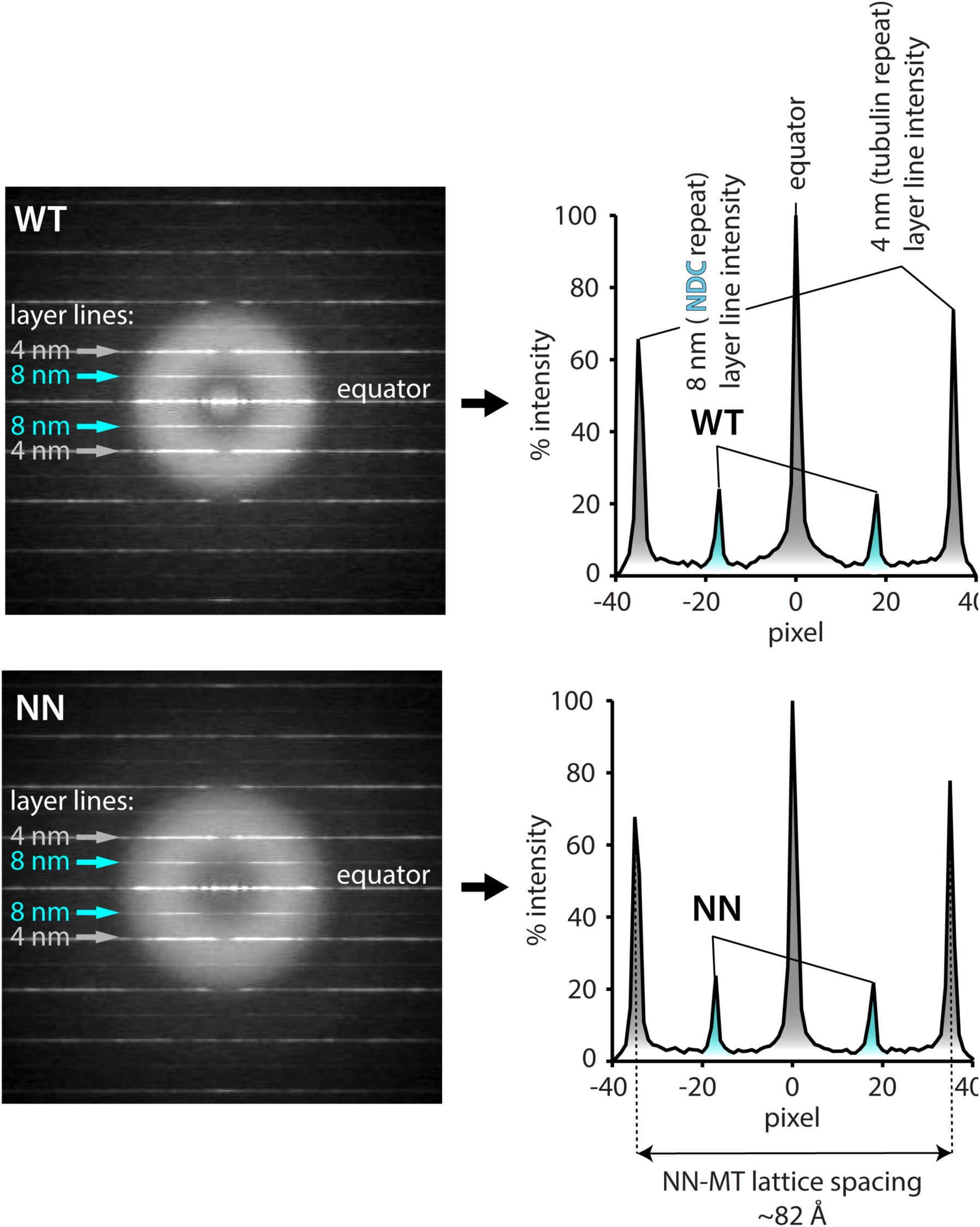
Fourier transformation of MT lattices decorated with WT or NN. Left: Averaged Fourier transforms of WT- and NN-decorated GDP-MT segments. Fourier transform of a DCX-MT image shows layer lines corresponding to 4 nm periodicity (tubulin subunits) and 8 nm periodicity (DC domain binding at MT lattice vertices). **Right**: Layer line intensities plotted using FiJi (fiji.sc). The plots are normalized against the equator intensity expressed as 100%. Relative MT lattice decoration by WT and NN can be assessed by the DC domain:tubulin layer line ratio. Similarly to WT-MT lattice, NN-MT lattice is compacted (∼82 Å spacing = tubulin repeat distance), as expected for the GDP state (Manka and Moores, 2018).

**Supplementary Fig. 9.**
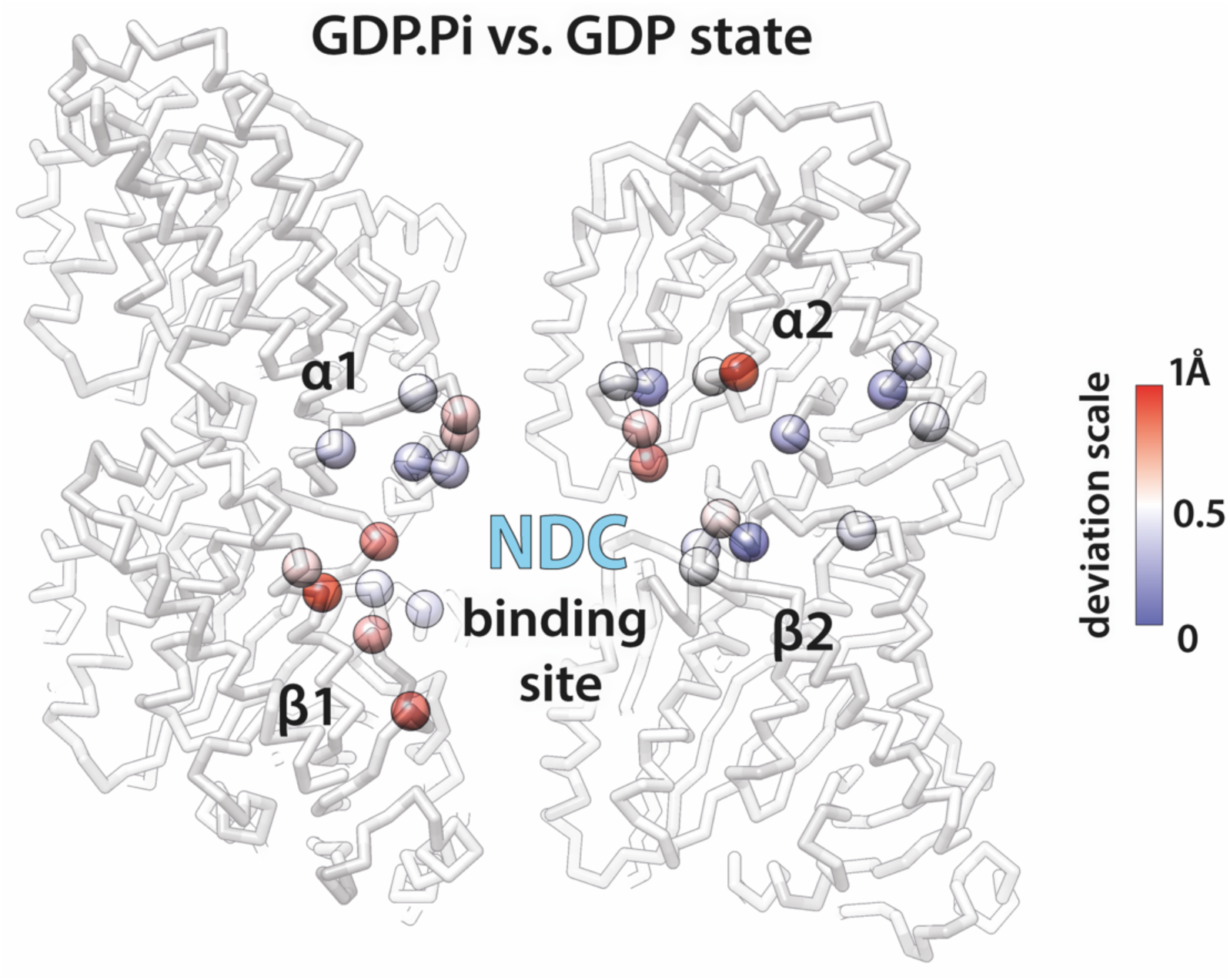
Differences between NDC binding sites in GDP.Pi- and GDP-MT lattices. Deviations in backbone (C*α*) atom positions are illustrated with colour-coded spheres.

**Supplementary Fig. 10.**
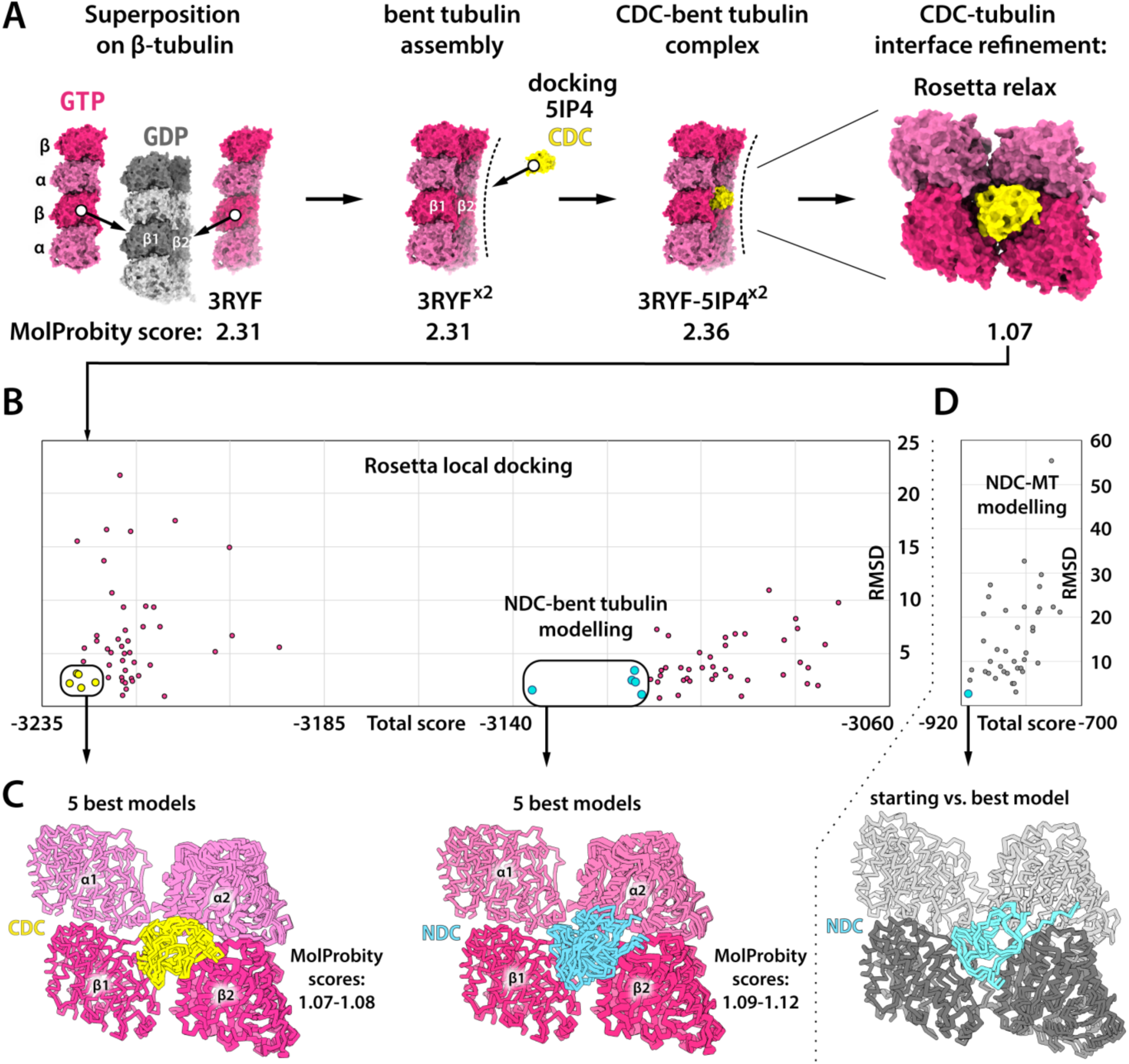
Modelling of CDC binding to curved tubulin assemblies. **A**. Building of CDC complex with bent tubulin assembly. X-ray structures of bent dimers of tubulin dimers (PDB: 3RYF) were aligned with an MT lattice fragment via *β*-tubulin subunits as indicated. Then an X-ray structure of CDC (PDB: 5IP4) was manually docked at the vertex of 4 bent tubulin dimers, within a 2-6 Å distance from each of the 4 tubulin subunits. The MolProbity score (Chen et al., 2010) of the popular structure validation server (http://molprobity.bio-chem.duke.edu/) combines the clashscore, rotamer and Ramachandran evaluations. Building of the complex did not introduce any major clashes as the score remains in good agreement with the original X-ray coordinates score. Next the CDC-tubulin interface was optimized using Rosetta relax protocol, improving the MolProbity score of the assembly to the value of 1.07. **B**. The refined complex of CDC with 4 tubulin subunits was subjected to full Rosetta local docking protocol to simulate 50 model structures. Total scores of these structures are plotted against RMSD. The same modelling and analysis were repeated using NDC instead of CDC. **C**. Superimposed backbones of the 5 models with the highest convergence (lowest RMSD and lowest total score). **D**. Control experiment. The full Rosetta local docking protocol (50 simulated models) applied to the cryo-EM NDC-MT structure from this study showed the expected convergence: best scoring model has the lowest RMSD from the experimental structure. The Rosetta total score scale is different from that shown in B due to different tubulin conformations - protein fold energy is a large component of the total score.

**Supplementary Fig. 11.**
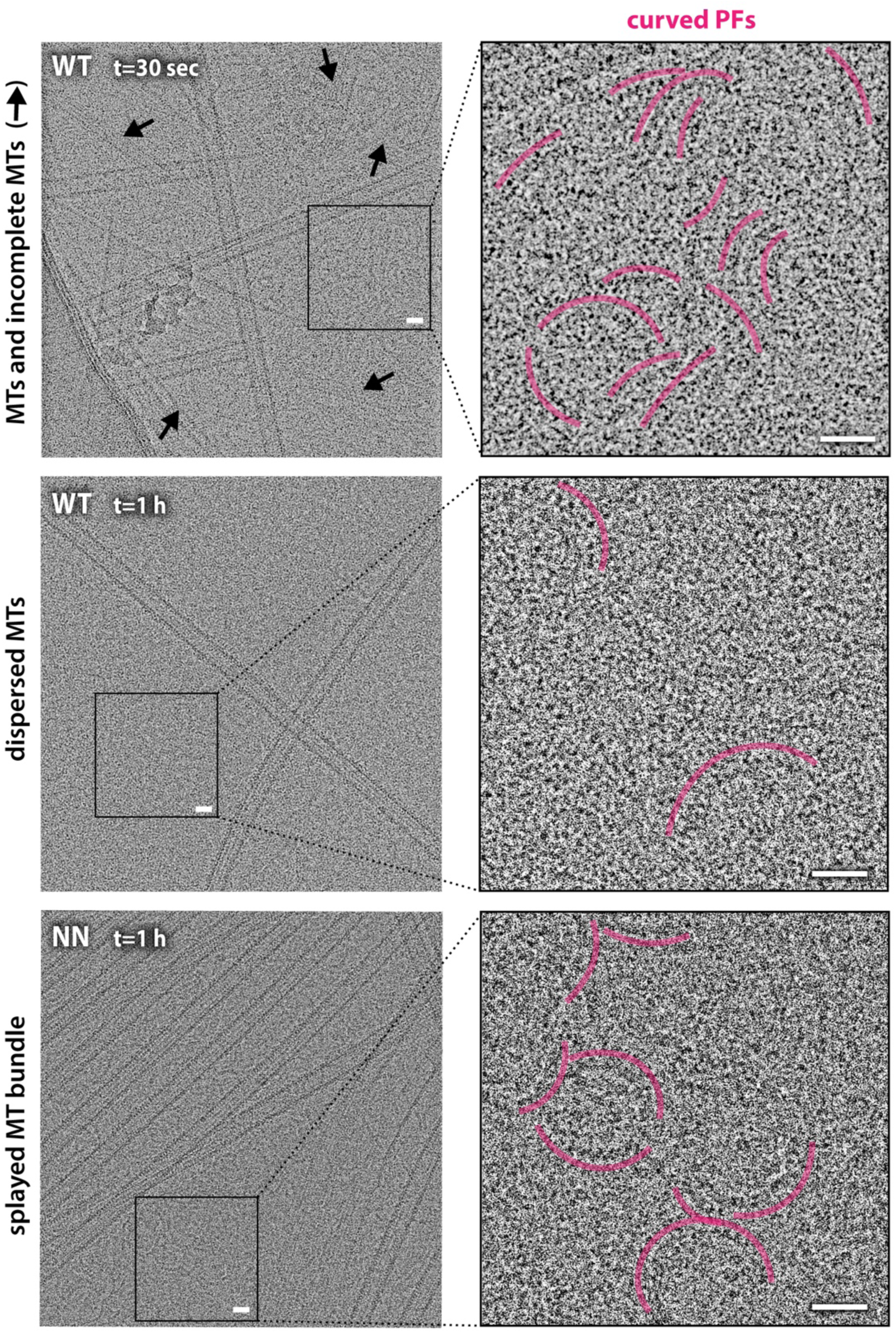
Tubulin polymerisation products in the presence of WT and NN. Left: Example cryo-EM micrographs showing WT-MTs and NN-MTs surrounded by other tubulin polymerisation products present at indicated polymerisation time points. **Right**: Close-up views of single and laterally associated non-MT curved tubulin protofilaments (PFs) showing various numbers and degrees of curvature, indicated with offset pink arches. Scale bar, 200 Å

**Supplementary Fig. 12.**
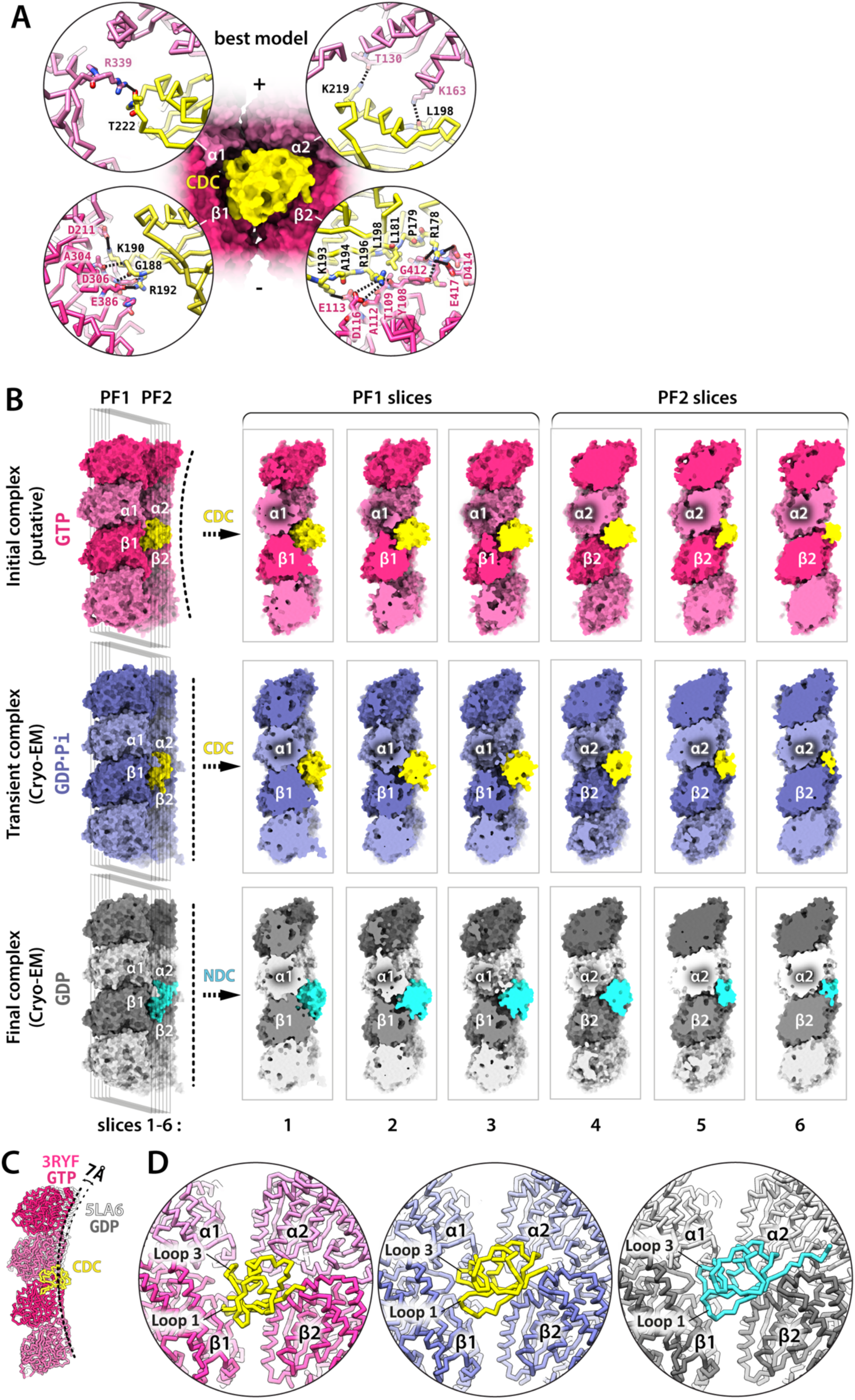
Comparisons of CDC-tubulin/MT and NDC-MT complexes. **A**. CDC-tubulin interaction sites in the highest-scoring model. Interacting residues are shown with sticks representation and coloured by heteroatom: O, red; N, navy blue. Likely polar contacts (<3.8 Å distance), solid lines; potential polar contacts (>3.8 Å distance), broken lines. **B**. Series of cross-sections through DC domain complexes with tubulin/MT designed to compare the fit between a DC domain and tubulin/MT at different stages of MT assembly. **C**. Alignment of X-ray structures of longitudinally associated bent tubulin dimers: 3RYF (GTP state; pink) and 5LA6 (GDP state; grey) shows different degrees of curvature between complexes of tubulin in different nucleotide states. CDC binding at the junction of tubulin assembly may further limit tubulin bending. **D**. Comparison of the DC domain backbone conformation at different stages of MT assembly. Coloured as in C.

**Supplementary Fig. 13.**
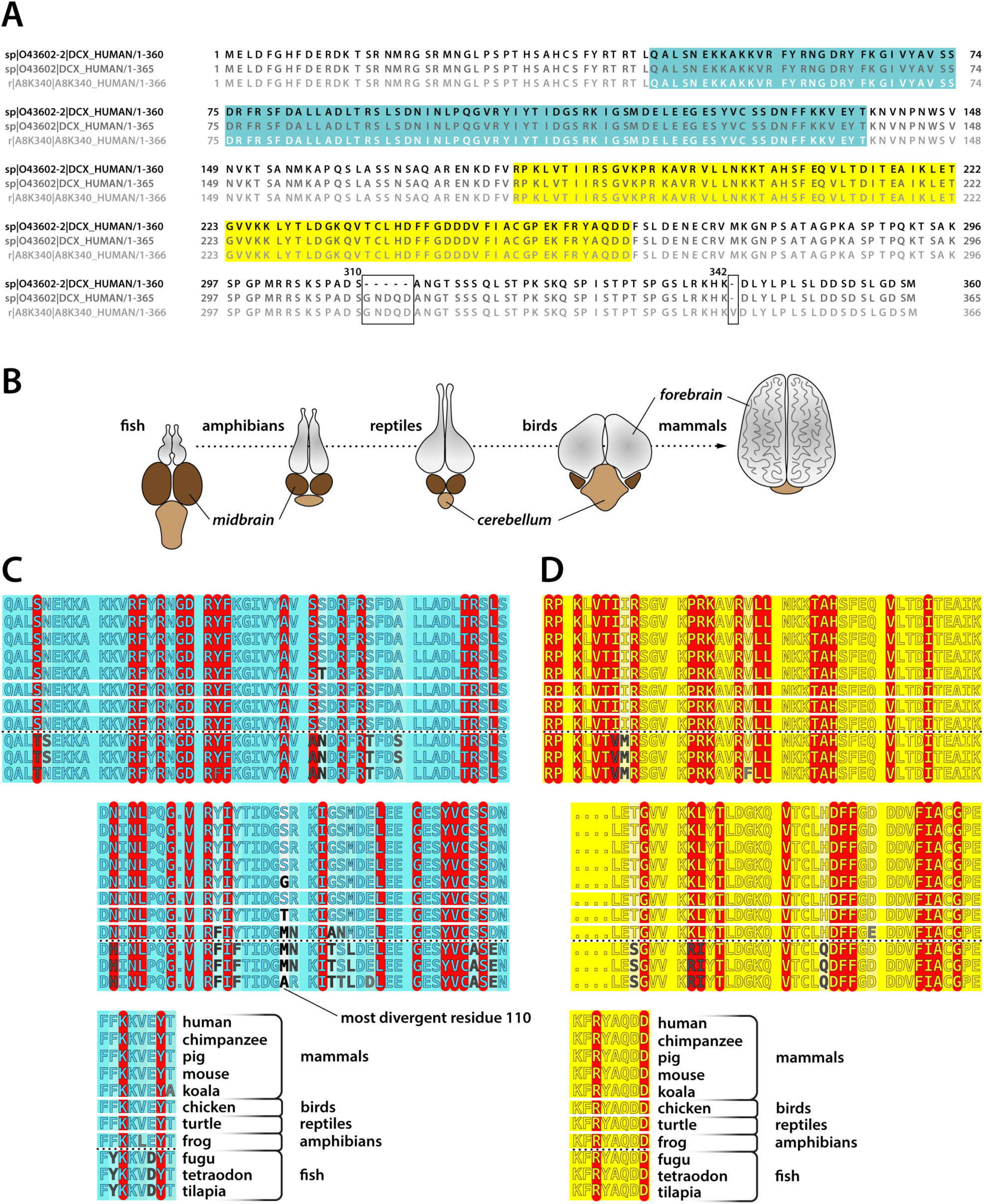
DCX isoforms in human and conservation of key DCX residues in different species. **A**. Three human splicing variants of DCX are aligned: 360 aa isoform 2 (this study; top sequence), 365 aa isoform 1 (middle sequence) and the 366 isoform (bottom). Sequence differences are boxed and NDC and CDC domain boundaries are indicated with the blue and yellow background, respectively. **B**. Schematic drawing of the major parts of the brain in different vertebrate classes, with an arrow indicating the relative forebrain growth. **C-D**. Alignments of NDC and CDC sequences, respectively, in selected vertebrate species. Residues identified as critically important for DCX function through the analysis human mutations data (Appendix Table S1) have been highlighted with red and divergent residues are marked with levels of grey, with most divergent sites coloured black. The degree of conservation is expressed with blue and yellow background intensity for NDC (in C) and CDC (in D), respectively: the fainter the colour the weaker the conservation. The most divergent site is residue 110 in NDC’s loop 4. It is one of the furthest residues from MT lattice in NDC, with a side chain projecting outward. Species selection includes model animals and an example outlier (koala) identified through large alignments of DCX sequences (>114 orthologs found in 183 species, including invertebrates) available through Ensembl genome database (ensembl.org). Koala was is one of the most divergent mammals regarding the DC domain sequences. Three fish species are shown to confirm fish class divergence (the only vertebrate class with dominant midbrain, as shown in A) from other vertebrate classes. Human, *Homo sapiens*; chimpanzee, *Pan troglodytes*; pig, *Sus scrofa*; mouse, Mus musculus; koala, Phascolarctos cinereus; chicken, Gallus gallus; turtle (painted turtle), *Chrysemys picta bellii*; frog, *Xenopus tropicalis*; fugu, *Takifugu rubripes*; tetraodon, *Tetraodon nigroviridis*; tilapia, *Oreochromis niloticus*.

**Supplementary Table 1.** Symptomatic missense mutations of doublecortin reported in humans.

| Mutation | Clinical significance | Condition | Review status | Last reviewed |
| --- | --- | --- | --- | --- |
| Pro64Leu* | LP | Lis | s | Jan 1 2018 |
| Met1Thr | P | H | s | Feb 8 2013 |
| Ser32Asn | P | H | s | Feb 8 2013 |
| Leu43Ser | P | H | s | Feb 8 2013 |
| Ser47Arg | P | Lis\|SLH | n.a. | Jan 9 1998 |
| Lys50Asn | P | H | s | Feb 8 2013 |
| Arg56Pro | LP | H | s | Apr 2 2014 |
| Phe57Cys | LP | H | s | Feb 8 2013 |
| Arg59Leu | P | Lis\|SLH | n.a. | Jan 9 1998 |
| Arg59His | P | H | s | Feb 8 2013 |
| Gly61Glu | P | H | s | Feb 8 2013 |
| Asp62Asn | P | Lis\|SLH | n.a. | Jan 9 1998 |
| Asp62Gly | LP | H | s | Feb 8 2013 |
| Arg63Cys | LP | H | s | Feb 8 2013 |
| Tyr64Cys | LP | H | s | Feb 8 2013 |
| Tyr145Asn* | LP | Lis | s | Sep 1 2017 |
| Phe65Leu | LP | H | s | Feb 8 2013 |
| Ala71Ser | P | Lis\|SLH | n.a. | Jan 28 2003 |
| Ser73Phe | P | H | s | Feb 8 2013 |
| Arg76Cys | LP | H | s | Apr 16 2013 |
| Arg78His | P | SLH | n.a. | Sep 1 1999 |
| Arg78Leu | P | H | s | Feb 8 2013 |
| Arg78Cys | P | H | s | Feb 8 2013 |
| Arg89Pro | P | H | s | Feb 8 2013 |
| Arg89Gly | P | SLH | n.a. | Sep 1 1999 |
| Leu91Pro | P | H | s | Feb 8 2013 |
| Asn94Asp | P | a.c.g. | s | Feb 8 2013 |
| Asn96Lys | P | F | s | Dec 3 2017 |
| Leu97Arg | P | H | s | Feb 8 2013 |
| Gly100Glu | LP | H | s | Feb 8 2013 |
| Gly100Val | P | H | s | Feb 8 2013 |
| Val101Met | P | H | s | Feb 8 2013 |
| Arg102Cys | P | H | s | Feb 8 2013 |
| Arg102His | P\|LP | H\|n.p. | m n.c. | Sep 30 2016 |
| Arg102Gly | P | n.p. | s | Jan 22 2016 |
| Ile104Phe | P | H | s | Feb 8 2013 |
| Ile113Phe | LP | H | s | Feb 8 2013 |
| Gly122Arg | P | H | s | Feb 8 2013 |
| Tyr125His | P | Lis\|SLH | n.a. | Jan 9 1998 |
| Val126Asp | LP | H | s | Feb 8 2013 |
| Cys127Tyr | LP | H | s | Feb 8 2013 |
| Ser129Leu | P | H | s | Aug 7 2013 |
| Tyr138His | P | H | s | Feb 8 2013 |
| Tyr138Cys | P | H | s | Feb 8 2013 |
| Lys151Glu | P | H | s | Feb 8 2013 |
| Lys174Glu | P | H | s | Feb 8 2013 |
| Arg178Cys | P | H | s | Feb 8 2013 |
| Arg178Gly | P | H | s | Feb 8 2013 |
| Arg178Leu | P | H | s | Feb 8 2013 |
| Lys180Glu | P | H | s | Feb 8 2013 |
| Val182Phe | P | Lis | s | Aug 23 2016 |
| Ile184Asn | LP | H | s | Feb 8 2013 |
| Arg186His | c.i. | Lis\|H | c.i. | Sep 1 2017 |
| Arg186Cys | P | H\|n.p. | m n.c. | Mar 30 2017 |
| Arg186Leu | P | H | s | Feb 8 2013 |
| Pro191Arg | P | H\|n.p. | m n.c. | May 30 2017 |
| Pro191Thr | LP | H | s | Feb 8 2013 |
| Arg192Trp | P | Lis\|SLH\|H\|n.p. | m n.c. | Dec 22 2016 |
| Arg196Ser | P | H | s | Feb 8 2013 |
| Arg196His | P | Lis\|SLH\|H | s | Feb 8 2013 |
| Arg196Gly | P | H | s | Feb 8 2013 |
| Arg196Cys | P\|LP | H\|n.p. | m n.c. | Jun 12 2017 |
| Leu199Pro | LP | H\|n.p. | m n.c. | Aug 8 2016 |
| Thr203Arg | P | Lis\|SLH | n.a. | Jan 9 1998 |
| Thr203Ala | P | H | s | Feb 8 2013 |
| Thr203Lys | P | H | s | Feb 8 2013 |
| Ala204Asp | P | H | s | Feb 8 2013 |
| His205Leu | LP | H | s | Feb 8 2013 |
| Val210Phe | P | H | s | Feb 8 2013 |
| Ile214Thr | P | H | s | Feb 8 2013 |
| Gly223Arg | P\|LP | H\|n.p. | m n.c. | Oct 13 2016 |
| Leu228Pro | P | H | s | Feb 8 2013 |
| Thr230Pro | LP | H | s | Feb 8 2013 |
| Phe242Ile | LP | H | s | Aug 27 2013 |
| Phe249Cys | LP | H | s | Feb 8 2013 |
| Ala251Ser | P | H | s | Feb 8 2013 |
| Ala251Asp | LP | n.p. | s | Jul 28 2015 |
| Gly253Arg | LP | n.p. | s | Jul 24 2017 |
| Arg258Pro | LP | H | s | Feb 8 2013 |

Mutations in grey lie outside of DC domains; a.c.g., abnormal cortical gyration; c.i., conflicting interpretations; F, fucosidosis; H, heterotopia; Lis, lissencephaly; LP, likely pathogenic; m, multiple submitters; n.a., no assertion; n.c., no conflicts; n.p., not provided; P, pathogenic; s, single submitter; SLH, subcortical laminar heterotopia; \*transcript variant 1, with 81 amino acids longer N-terminal sequence (starting with Met -81 with respect to transcript variant 2 relevant in this study), Pro64 in transcript 1 does not exist in DCX isoform from this study and Tyr145 is equivalent with Tyr64 in DCX isoform from this study.

**Movie 1. Morphing of the transition of the CDC-bound bent GTP-tubulin assembly model to the CDC-bound GDP.Pi-MT structure.** The structures are in backbone representations. Bent GTP-tubulin, pink; straight GDP.Pi-tubulin, violet; CDC, yellow.

